# Cell-based multisensor array for detecting cancer biomarkers in human urine

**DOI:** 10.1101/2025.07.18.665501

**Authors:** Hisatoshi Mimura, Toshihisa Osaki, Haruka Oda, Sho Takamori, Yuki Kodama, Norio Sasaoka, Yasuhiko Takahashi, Shoji Takeuchi

## Abstract

Detecting volatile urinary biomarkers offers a compelling strategy for noninvasive diagnosis of diseases such as cancer. Biohybrid sensors employing sensor cells that express olfactory receptors—with finely tuned molecular recognition capabilities—are garnering increasing attention. However, their application to physiological samples remains challenging due to variability in sensor cell responsiveness and limited signal intensity. Here, we report a volatile biomarker detection platform based on a multisensor array in which insect olfactory receptor– expressing cells are encapsulated in hydrogel and housed within microwells featuring vertical slits. This structural design enables dense three-dimensional cell immobilization and rapid, uniform exposure to target molecules, enhancing the sensitivity of fluorescence responses and minimizing signal variability. Multiple types of sensor cells on a single array enabled the selective and simultaneous detection of potential cancer-related target molecules—acetophenone, phenol, and 6-methyl-5-hepten-2-one. Furthermore, through hexane extraction combined with gas-phase delivery, we reliably detected acetophenone introduced into human urine at micromolar concentrations. This platform offers a compact, reproducible, and scalable solution for volatile biomarker detection using cell-based sensors.

## Introduction

Biomarkers in human urine, a non-invasively collectable biofluid, have been proposed as potential candidates for early cancer detection [1–3]. Developing sensor technologies with high sensitivity and selectivity for detecting these biomarkers is essential [4, 5]. Biohybrid sensors utilizing the excellent molecular recognition capabilities of biological systems are gaining attention as an emerging approach [6–12]. Olfactory receptors (ORs), expressed on the cell surface, function as primary sensors in biological olfaction, recognizing target molecules with high specificity and sensitivity [13, 14].

Biohybrid sensors utilizing ORs can be categorized into cell-free [15, 16] and cell-based types [17–20]. In cell-free sensors, purified ORs are either used as isolated proteins or incorporated into liposomes or nanodiscs, and then immobilized onto transducers to function as molecular recognition elements for capturing target molecules [21–35]. Additionally, ORs can be reconstituted into lipid bilayers, where their electrophysiological responses upon ligand binding are recorded [15, 16]. On the other hand, cell-based sensors detect intracellular responses triggered by target molecules. These responses can be detected as changes in fluorescence intensity (Fig. 1A), as well as through transducer-based and electrophysiological methods [17–19, 36–42].

**Fig. 1.**
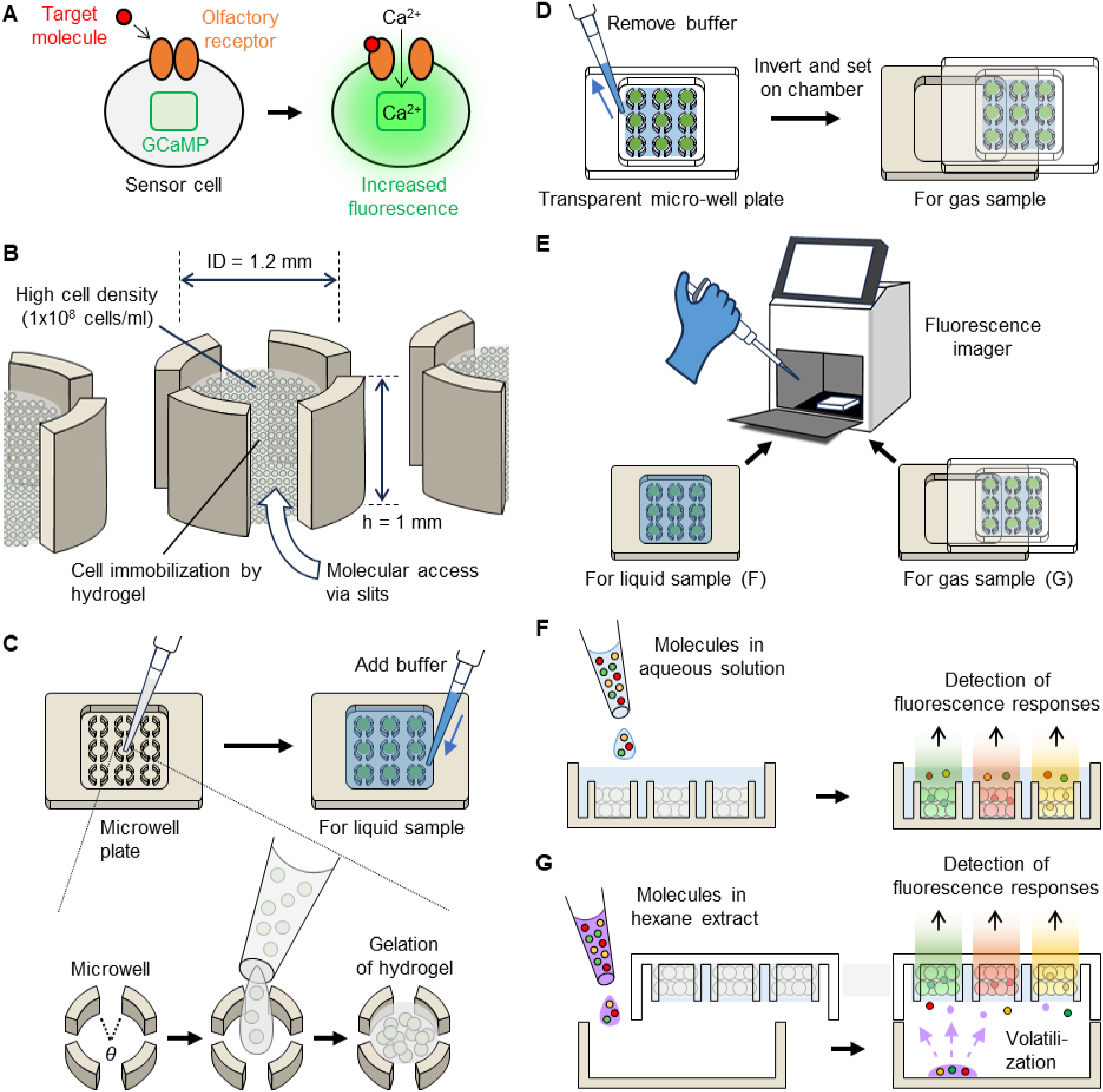
Overview of the proposed sensor cell array system for detecting target molecules. (**A**) Schematic of a sensor cell responsive to a target molecule. (**B**) Structure of slit-integrated microwells enabling hydrogel-based immobilization of densely packed sensor cells (>10⁵ cells/well) and efficient molecular access. (**C**) Fabrication of the cell array in microwells using pipetting. (**D**) Setup for gas-phase detection using the array. (**E**) Measurement procedures for liquid and gas samples using a fluorescence imager. (**F**, **G**) Fluorescence responses of multisensor arrays to target molecules in liquid samples (F) and in hexane-extracted gas samples (G).

A key advantage of cell-based sensors lies in eliminating the need for complex purification processes of ORs expressed in cells or cell-free translation systems, enabling the stable utilization of multiple OR types [8, 9, 11]. Olfactory cells and genetically engineered cultured cells that express ORs—collectively referred to as sensor cells—have been extensively studied for their application in target molecule detection [8, 15]. In recent years, technologies have been developed to array multiple types of sensor cells on a small substrate for the detection of multiple target molecules [17, 37, 39, 41, 43]. Furthermore, techniques for detecting target molecules in the gas phase using sensor cells have also been developed [18, 38, 42, 44–47]. However, cell-based sensors have yet to be applied for biomarker detection in physiological samples. This limitation primarily arises from the inherent variability in individual sensor cell responses and the weak signal intensity of cellular outputs [47], which together make it challenging to achieve sufficient sensitivity for biomarker detection. Therefore, the development of small, easy-to-use, and highly sensitive detection platform—suitable for point-of-care diagnostics—is essential for the practical implementation of cell-based sensors.

In this study, we develop a cell-based sensor platform capable of detecting biomarkers by leveraging fluorescence signal responses from high-density sensor cells (Fig. 1B). By capturing the cumulative fluorescence response from a densely packed sensor cell population, we mitigate signal variability among individual cells and enhance detection sensitivity. For this purpose, we introduce a microwell with slits on its sidewalls, enabling the efficient arraying of hydrogel-encapsulated, high-density sensor cells (Fig. 1C). This microwell design additionally improves target molecule accessibility, facilitates the activation of a large population of sensor cells, and further enhances detection sensitivity (Fig. 1B). The plate, composed of multiple microwells within a single chamber, also allows for stable measurements and efficient parallel analysis using a fluorescence imager, supporting the detection of multiple target molecules (Fig. 1C, E, F). Here, we investigate the response characteristics of the sensor cell array fabricated using the microwell plate, including its selectivity and sensitivity, and evaluate its multiplex detection capability in mixtures of target molecules. As a practical application, we demonstrate the detection of a potential cancer biomarker, extracted from human urine with a volatile organic solvent, in the gas phase using the developed sensor (Fig. 1D, E, G).

## Results

### Hydrogel-based immobilization of sensor cells in slit-integrated microwells

To evaluate the reliable immobilization of sensor cells encapsulated in hydrogel, we fabricated slit-integrated microwells formed by partial arc-shaped microprotrusions (Fig. 1B). Each microwell had a cylindrical shape with an inner diameter of 1.2 mm, a height of 1.0 mm, and an internal volume of approximately 1.1 μL.

The sensor cells used in this study were cultured insect cells that stably expressed an olfactory receptor (OR) and a co-receptor (Orco) from insects [48, 49] and a calcium-sensitive fluorescent protein (jGCaMP7) [50] (Fig. 1A). When target molecules bind to the OR complex on the cell membrane, the intrinsic ion channel opens, allowing calcium ions to flow into the cytoplasm and thereby increasing GCaMP fluorescence. The stable cell lines were generated in this study, and the detailed method is described in the Materials and Methods section. In the initial experiments, we used sensor cells expressing the *Aedes aegypti*-derived olfactory receptor AaOR15, which specifically recognizes acetophenone [51]—a potential biomarker for lung and breast cancers [52, 53]. AaOR15 was selected as a representative receptor for validating sensor function in the microwells. In subsequent experiments using a multi-sensor array, we employed cells expressing different ORs to enable multiplexed odorant detection.

First, we assessed the sample-holding capability of the slit-integrated microwells. As shown in Fig. 2A, Milli-Q water was dispensed into a microwell using a micropipette, and the droplet was retained within the microwell without leakage by surface tension. Furthermore, Movie 1 shows that continuous dispensing using an electric micropipette could be achieved with stable operation. The dispensed droplets remained confined within individual microwells without fusing with neighboring wells (Supplementary Fig. 1A, B), suggesting that sensor cell–hydrogel mixtures could also be similarly dispensed without cross-contamination. Next, to examine whether the system could accommodate large-scale dispensing, we fabricated a microwell plate composed of 81 slit-integrated microwells arranged in a 9×9 format (Fig. 2B; Supplementary Fig. 2D, H). When colored Milli-Q water was dispensed, solutions with different concentrations were independently placed into each microwell (Fig. 2C).

**Fig. 2.**
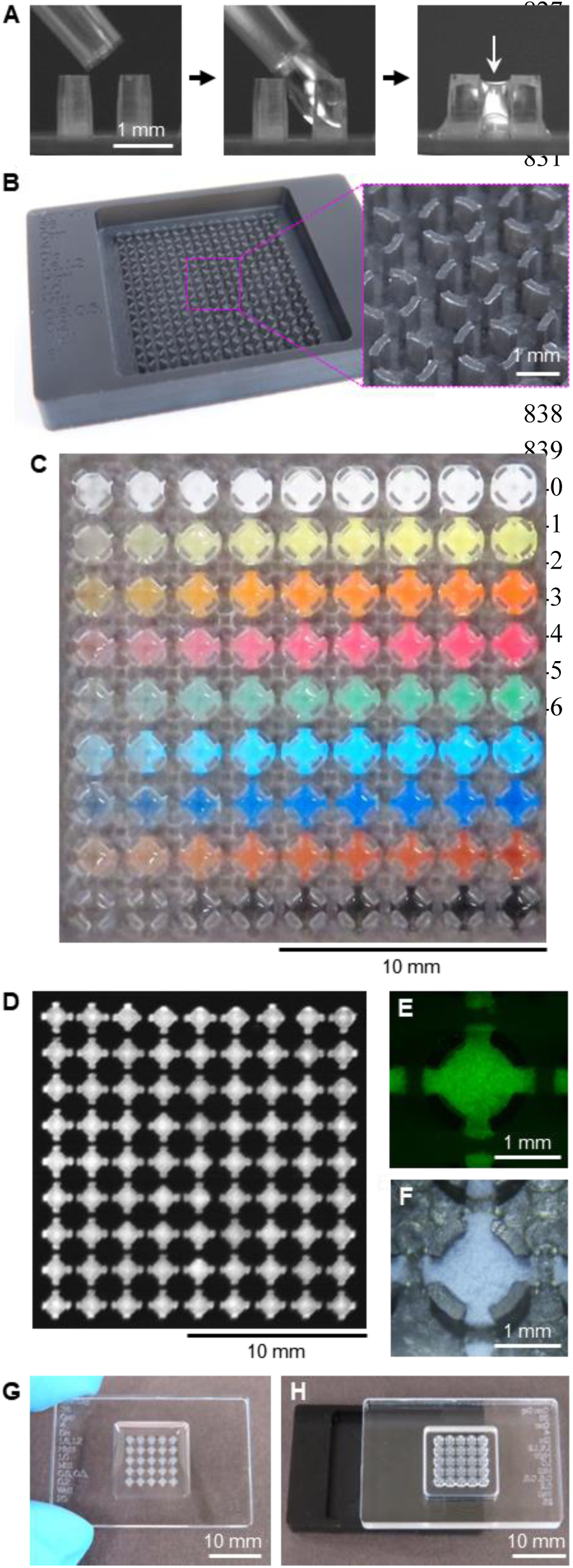
Arraying of sensor cells using a slit-integrated microwell plate. (**A**) Side-view photographs showing the dispensing of 1 μL of Milli-Q water into a slit-integrated microwell formed by acrylic microstructures. The white arrow indicates a stably retained water droplet. (**B**) Photograph of a fabricated slit-integrated 9×9 microwell plate made of black acrylic. (**C**) Gradient pattern generated by dispensing water-based paint–colored Milli-Q water into a slit-integrated 9×9 microwell plate made of transparent acrylic. (**D**) Fluorescence image of same-type sensor cells arrayed in the 9×9 microwell plate. (**E**, **F**) Fluorescence (E) and bright-field (F) microscopic images of hydrogel-immobilized sensor cells in a single microwell. (**G**) Sensor cell array prior to buffer removal. (**H**) Measurement setup for gas-phase detection using the inverted array shown in (G) after buffer removal.

Using this system, we mixed the AaOR15-expressing sensor cells with a liquid hydrogel precursor and dispensed the mixture into the 9×9 microwell plate. Upon gelation, the hydrogel successfully immobilized the sensor cells within each microwell (Fig. 2D). Microscopic observations confirmed that the sensor cells were immobilized within the microwells and encapsulated in hydrogel, as shown by fluorescence and bright-field imaging (Fig. 2E, F). These results demonstrate that the slit-integrated microwell plate enables simple and reliable fabrication of hydrogel-encapsulated sensor cell arrays by pipetting.

### Effect of sensor cell density and microwell slit structure on fluorescence response sensitivity

#### Sensor cell density

To investigate the effect of sensor cell density on fluorescence response sensitivity, hydrogel-encapsulated sensor cell arrays were fabricated (Fig. 1C), and their responses to target molecules in aqueous samples were measured using a fluorescence imager (Fig. 1E). A 3×3-format plate containing 9 microwells was used for the experiments (Supplementary Fig. 2A, E).

Figure 3A shows pairs of pseudo-colored fluorescence images of the AaOR15-expressing sensor cell arrays fabricated at cell densities of 1×10⁶ and 1×10⁸ cells/mL before and after the addition of acetophenone. The nine pseudo-colored fluorescence spots in each array correspond to microwells where the hydrogel-encapsulated sensor cells were immobilized. Upon the addition of acetophenone (final concentration: 100 μM), little fluorescence change was observed at 1×10⁶ cells/mL, whereas a clear increase was detected at 1×10⁸ cells/mL. The time-course plot of the fluorescence response, expressed as the normalized change in fluorescence intensity (ΔF/F₀; see Methods for details), demonstrated that fluorescence responses increased in a cell density-dependent manner between 1×10⁶ and 1×10⁸ cells/mL Fig. 3B). Furthermore, comparison of the maximum value of the normalized fluorescence response (Max ΔF/F₀) and the coefficient of variation (CV) revealed that the maximum response increased more than 15-fold and the CV decreased to less than half at 1×10⁸ cells/mL compared to 1×10⁶ cells/mL (Fig. 3C).

**Fig. 3.**
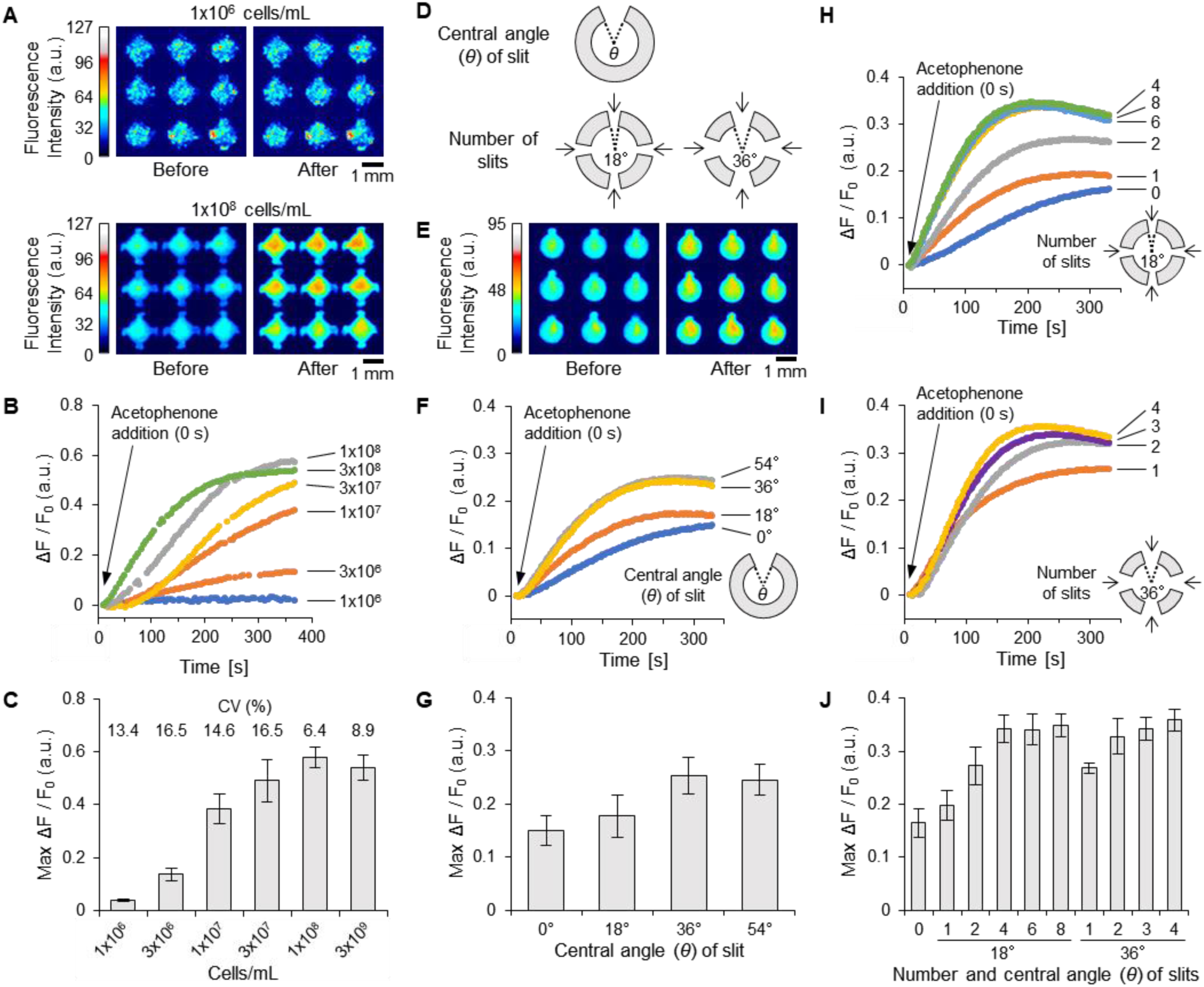
Effects of sensor cell density and slit structures in microwells on the fluorescence responses of sensor cell arrays. (**A**) Pseudocolor fluorescence images showing responses to the target molecule in sensor cell arrays with different cell densities (1×10⁶ and 1×10⁸ cells/mL). Before: pre-response (11 s after addition); After: post-response (300 s after addition). (**B**) Time-course fluorescence responses at various cell densities (1×10⁶ to 1×10⁸ cells/mL). (**C**) Maximum responses from (B), with coefficients of variation (CV) indicated above each bar. (**D**) Schematic of slit structures defined by central angle (θ); both single and multiple slits (θ = 18° or 36°) are shown. (**E**) Pseudocolor fluorescence images showing responses using microwells with a single 36° slit. Before: 11 s; After: 300 s after addition. (**F**) Time-course fluorescence responses using microwells with no slit (θ = 0°) or a single slit (θ = 18°, 36°, or 54°). (**G**) Maximum responses from (F). (**H**, **I**) Time-course fluorescence responses using microwells with multiple slits: (H) θ = 18° (0–8 slits); (I) θ = 36° (1–4 slits). (**J**) Maximum responses from (H) and (I). Sensor cells expressing AaOR15 were used, with acetophenone (100 μM final) as the target molecule. Each curve represents the mean of *n* = 9; each bar shows mean ± SD, *n* = 9. Buffers used were HBSS/HEPES (pH 7.2). Excitation was intermittent (2 s every ∼5 s) in (A–C), and continuous in (E–J). Note: Higher fluorescence responses in (A–C) compared to (E–J) are likely due to reduced photobleaching under intermittent excitation.

These results indicate that high-density packing of sensor cells within microwells is crucial for enhancing fluorescence sensitivity and reducing variability. The increase in normalized fluorescence response can be explained by the suppression of background signal dilution, as densely packed cells produce a stronger cumulative signal. The decrease in CV results from a stable increase in fluorescence intensity with a consistent standard deviation (SD), which is likely due to both suppression of cell detachment by hydrogel encapsulation and improved uniformity of cell distribution within the microwell through high-density packing. Note that the decrease in fluorescence response at 3×10⁸ cells/mL was due to the collapse of the hydrogel structure.

A simple calculation suggests that filling a microwell with a diameter of 1.2 mm and a height of 1 mm with 1.1 μL of a 1×10⁸ cells/mL suspension results in approximately 50 layers of cells with a diameter of about 20 μm. Each layer contains approximately 2200 cells, corresponding to 2000 cells/mm². By observing more than two layers simultaneously using an optical system with a long depth of field, fluorescence detection becomes feasible at cell densities that exceed the theoretical maximum density of hexagonal close packing in two dimensions (packing fraction of 90.7%, approximately 3200 cells/mm²). Furthermore, the experimentally optimized cell density of 1×10⁸ cells/mL approaches the theoretical limit for hexagonal close packing in three dimensions (maximum packing fraction of 74%), corresponding to approximately 1.8×10⁸ cells/mL. These findings indicate that the selected cell density provides an appropriate balance between fluorescence sensitivity and hydrogel structural stability. Based on these results, a sensor cell density of 1×10⁸ cells/mL was adopted for subsequent experiments.

#### Microwell slit structure

To assess the effect of slit structure on fluorescence response, microwells with a single slit of varying central angles (θ = 0° [no slit], 18°, 36°, 54°, 72°, or 90°) were fabricated. The slit width was defined by the central angle θ (Fig. 3D), and the hydrogel retention capabilities of these microwells were evaluated (Supplementary Fig. 3). Microwells with slits of θ ≥ 54° exhibited increased variability in hydrogel retention due to leakage through the slits. Therefore, fluorescence responses were evaluated using microwells with slit angles ranging from 0° to 54°.

As shown in Fig. 3E, a AaOR15-expressing sensor cell array using microwells with a single 36° slit exhibited a clear fluorescence increase upon acetophenone addition. Time-course analysis revealed that fluorescence responses began approximately 20 seconds after acetophenone addition and reached a maximum around 250 seconds (Fig. 3F). Comparison of maximum fluorescence responses showed that microwells with a single slit began to exhibit enhanced responses at θ = 18°, reaching maximum enhancement at θ = 36°, with approximately 1.7-fold improvement compared to microwells without slits (Fig. 3G).

After identifying the optimal slit angle, the effect of the number of slits was then investigated. Microwells containing 1–8 slits (18°) or 1–4 slits (36°) were fabricated (Fig. 3D; Supplementary Fig. 4). Time-course data showed that increasing the number of slits shortened the time to reach the maximum fluorescence response, with microwells having four or more slits achieving maximum responses approximately 20 seconds earlier than those with a single slit (Fig. 3H, I). As shown in Fig. 3J, the maximum fluorescence responses were comparable between microwells with four or more slits for both the 18° and 36° conditions, and were approximately 2.1-fold higher than those observed in microwells without slits.

The fluorescence responses of hydrogel-encapsulated sensor cells immobilized in microwells were enhanced by increasing the slit opening area and number. However, responses were maximized with four or more slits of either 18° or 36°, and further increases in slit number did not yield additional enhancement. Similar fluorescence responses were obtained under different opening areas (2.64 mm²: 18°×8 or 36°×4 slits, and 1.88 mm²: 18°×4 slits), suggesting that proper spatial arrangement of slits, rather than simply expanding the opening area, is critical for maximizing sensitivity. These findings indicate that slits facilitate multidirectional diffusion of target molecules into the hydrogel, leading to more uniform and rapid exposure of sensor cells and promoting simultaneous responses. Under the present experimental conditions, a four-slit configuration was sufficient to fully harness sensor cell performance. Considering fabrication advantages, such as easier machining and shorter processing time using larger-diameter end mills, microwells with four 36° slits were adopted for subsequent experiments.

### Effect of sample addition on sensor cell response reproducibility

To evaluate the effect of the sample addition method on response consistency across microwells, hydrogel-encapsulated sensor cell arrays were fabricated using a 9×9 microwell plate (Fig. 2B; Supplementary Fig. 2D, H).

Figures 4A and 4B show fluorescence images of AaOR15-expressing sensor cell arrays before and after the addition of acetophenone solution, either by a single addition near the center of the plate (Fig. 4A) or by four separate additions at peripheral sites (Fig. 4B). In the single addition case, fluorescence responses were absent in some microwells, and the time-course fluorescence signals exhibited considerable variability (Fig. 4C). In contrast, separate additions at four peripheral points resulted in a more uniform fluorescence response across the entire array and more stable time-course behavior (Fig. 4D).

**Fig. 4.**
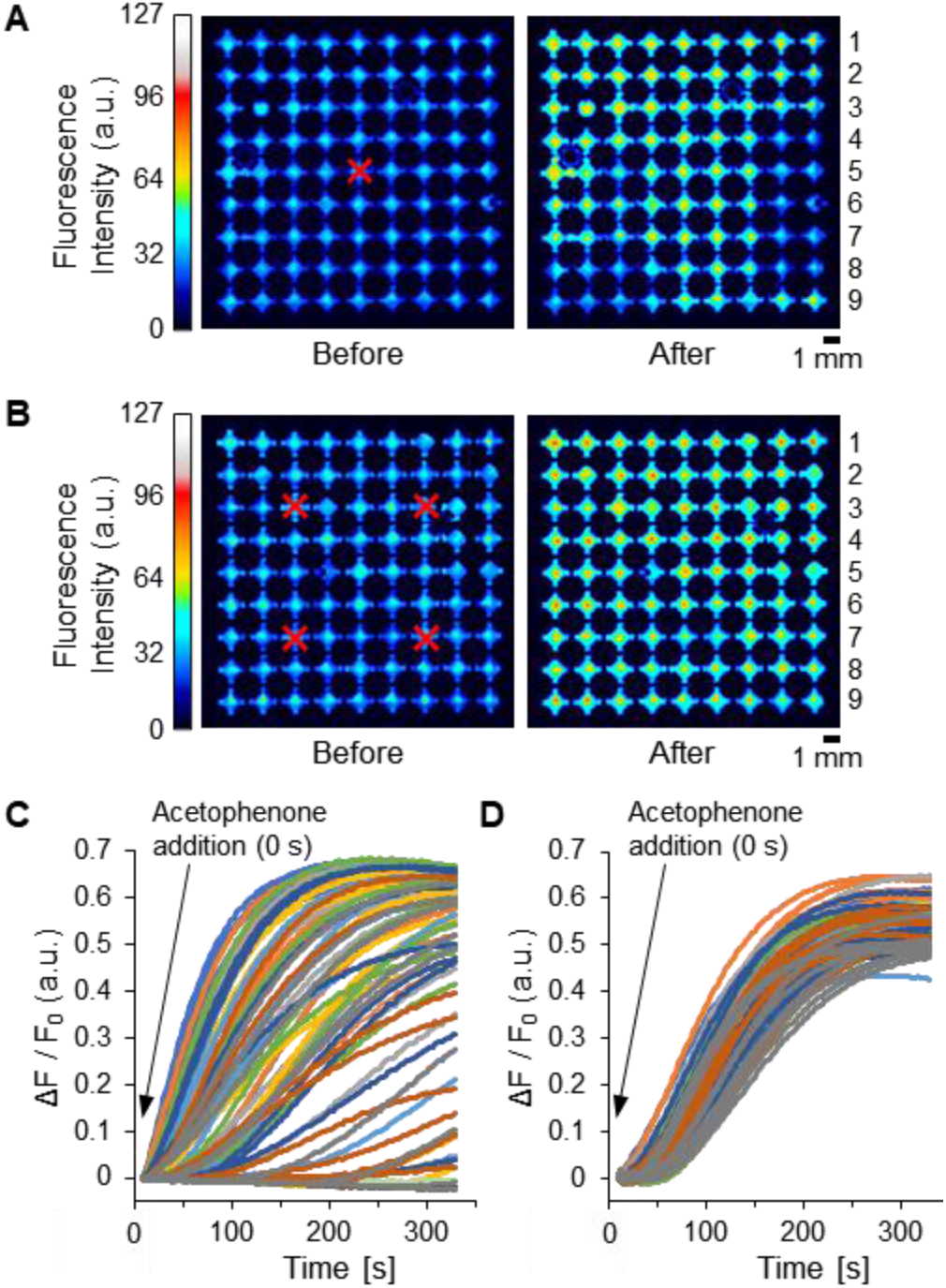
Effects of the number of target molecule additions on the consistency of sensor cell array responses. (**A**, **B**) Pseudocolor fluorescence images showing the responses of a sensor cell array composed of AaOR15-expressing cells in a slit-integrated 9×9 microwell plate, each microwell containing four slits (central angle = 36°). Red crosses indicate where equal volumes of acetophenone solution (100 μM final) were added either in a single step (A) or in four separate steps (B). Before: pre-response (9 s after addition in A; 12 s in B); After: post-response (300 s in both). (**C**, **D**) Time-course fluorescence responses corresponding to (A) and (B), for single (C) and four separate (D) additions, respectively. Line colors indicate microwell plate rows: blue (row 1), orange (2), gray (3), yellow (4), light blue (5), green (6), dark blue (7), dark orange (8), dark gray (9).

Further evaluations were conducted using 3×3, 5×5, and 7×7 microwell plates (Supplementary Fig. 2). In all formats, separate additions reduced the variability of sensor cell array responses compared to single additions, with particularly pronounced improvements in the 7×7 and 9×9 formats (Fig. 4; Supplementary Fig. 5). The coefficient of variation (CV) of the maximum fluorescence response increased with array size under single addition conditions, whereas the CV remained consistently low across all array sizes under separate addition conditions (Supplementary Fig. 6). Specifically, the CVs under single addition were 9.6% (3×3), 13.4% (5×5), 40.5% (7×7), and 48.8% (9×9). In contrast, under separate addition, the CVs were 9.2%, 5.3%, 5.7%, and 8.0% for these array formats, respectively. These findings demonstrate that achieving uniform exposure to target molecules is critical for enhancing the consistency of sensor cell array responses. Combined with an appropriate sample addition strategy, the developed microwell plate provides a reliable platform for precise multisensor array detection.

### Fabrication of multisensor arrays and characterization of responses to target molecules

To evaluate the responses of a multisensor array to target molecules, nine types of sensor cells were arrayed, and their selectivity toward three target molecules was examined. The sensor cells included those expressing AaOR15, as well as AgOR1, AgOR2, AgOR10, AgOR11, AgOR21, and AgOR46 (derived from *Anopheles gambiae*), AaOR4 (derived from *Aedes aegypti*), and DmOR47 (derived from *Drosophila melanogaster*) [51, 54–56]. In addition to acetophenone (described above), we also used phenol and 6-methyl-5-hepten-2-one (6M5H2O) as target odorants. Phenol has been reported as a potential biomarker for lung, thyroid, gastric, and esophageal cancers [3, 57–59], while 6M5H2O has been specifically linked to gastric cancer [60–62].

Multisensor arrays were fabricated by arranging nine microwells per sensor cell type on a 9×9 microwell plate. Fluorescence responses were measured before and after the addition of target molecule solutions using four sequential pipetting operations (Fig. 5A; Supplementary Fig. 7A). Variations in initial fluorescence intensity among sensor cell types, despite constant cell concentrations (1×10⁸ cells/mL), suggest intrinsic differences in the expression levels of the calcium sensitive fluorescent protein across the stably expressing cell lines. Upon acetophenone addition, fluorescence responses varied depending on the sensor cell type, with the highest response observed in AaOR15-expressing cells (Fig. 5B). Comparisons with the negative control (DMSO) and with other target molecules (phenol or 6M5H2O) revealed distinct response patterns across sensor cell types (Supplementary Fig. 7). As shown in Fig. 5C–F, none of the sensor cells responded to the negative control, while several exhibited selective responses to specific target molecules. In particular, AgOR1, AaOR4, and AaOR15 showed high selectivity to phenol, 6M5H2O, and acetophenone, respectively, consistent with previously reported characteristics [51, 54, 55]. These results indicate that the multisensor arrays fabricated using the microwell plate can be effectively applied for the selective detection of target molecules.

**Fig. 5.**
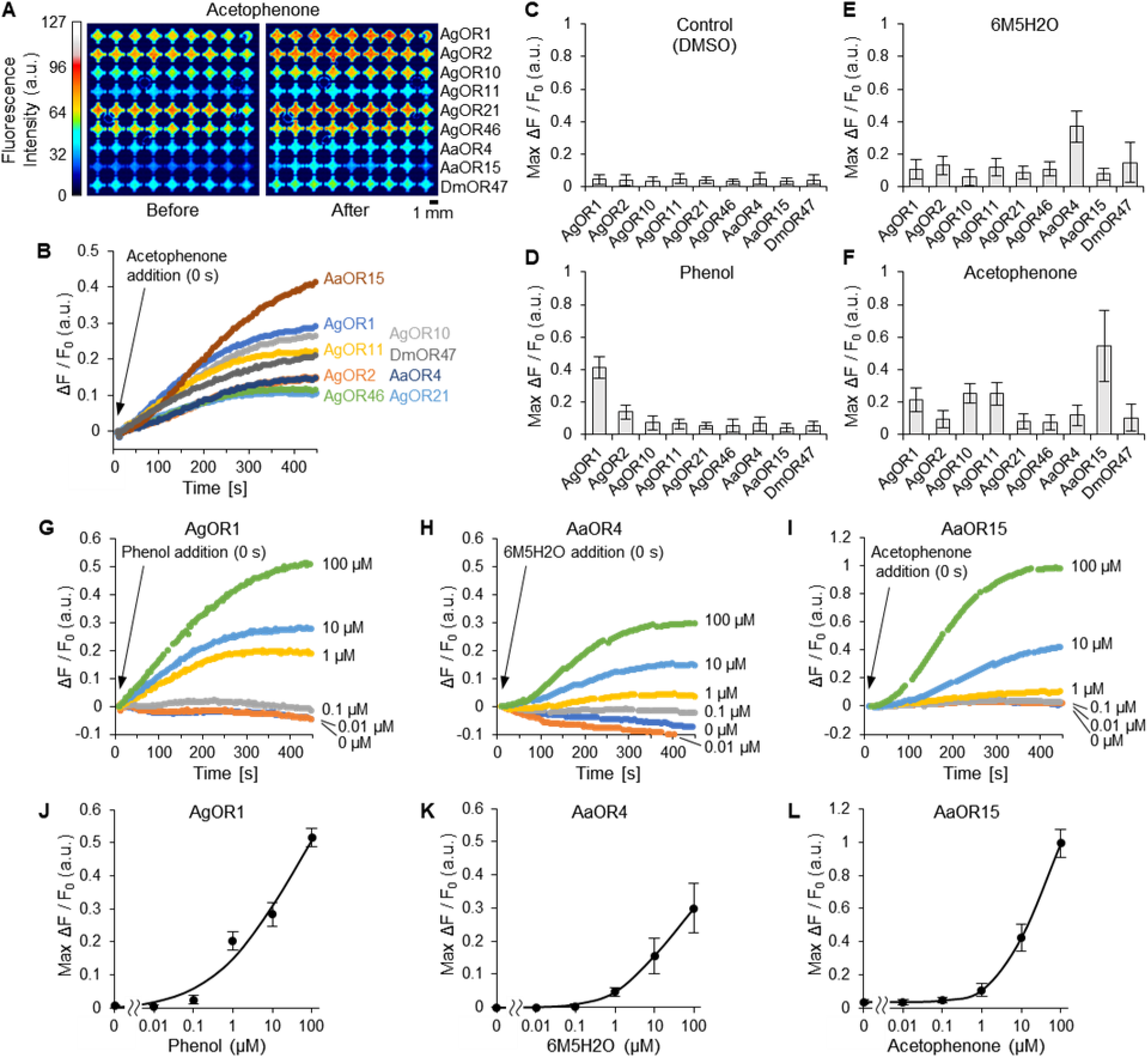
Selectivity and dose-dependent responses of sensor cells using sensor cell arrays. (**A**) Pseudocolor fluorescence images showing the responses of a multisensor array composedf nine sensor cell types, each expressing a different olfactory receptor (OR) and arranged in horizontal rows by OR type. Acetophenone (100 μM final) was added in four separate steps. Before: pre-response (11 s after addition); After: post-response (401 s after addition). (**B**) Time-course fluorescence responses under the same conditions as in (A). Each curve represents the mean of *n* = 9. (**C**–**F**) Maximum fluorescence responses of the nine sensor cell types to control (DMSO, no target molecule) (C), phenol (D), 6-methyl-5-hepten-2-one (6M5H2O) (E), and acetophenone (F). All target molecules were used at 100 μM in 0.1% DMSO. Bars represent mean ± SD; *n* = 36 (DMSO, 6M5H2O, acetophenone), *n* = 27 (phenol). (**G**–**I**) Time-course fluorescence responses of sensor cells expressing AgOR1 (G), AaOR4 (H), and AaOR15 (I) to increasing concentrations of their respective target molecules. Each cell type was arrayed in a slit-integrated 3×3 microwell plate. Curves represent the mean of *n* = 9. (**J**–**L**) Dose–response curves showing the maximum fluorescence responses from (G–I). Each data point represents the mean ± SD; *n* = 9.

Further evaluations were conducted on AgOR1-, AaOR4-, and AaOR15-expressing sensor cells, which demonstrated high selectivity, to assess their response sensitivities in more detail. Sensor cell arrays were fabricated using 3×3 plates for each sensor type, and fluorescence responses were measured upon exposure to target molecule solutions of varying concentrations. As shown in Fig. 5G–I, fluorescence responses increased with increasing target molecule concentrations.

Moreover, plots of the maximum fluorescence responses against the logarithm of target molecule concentrations (Fig. 5J–L) clarified the dynamic range of fluorescence responses for each sensor cell array. The limit of detection (LOD) was defined as the lowest tested concentration at which the maximum fluorescence response exceeded the baseline fluorescence level plus three times its SD. The baseline was set at 0 μM for phenol and acetophenone, and 0.01 μM for 6M5H2O. Based on this definition, among the concentrations tested in this study, the LODs for phenol, 6M5H2O, and acetophenone were 0.1, 1, and 1 μM, respectively, using sensor cell arrays expressing AgOR1, AaOR4, and AaOR15. The obtained detection sensitivities were consistent with the previously reported response ranges for each olfactory receptor [51, 55, 63].

### Simultaneous detection of multiple target molecules using a multisensor array

To evaluate whether the developed multisensor array can simultaneously and selectively detect multiple target molecules, a sensor cell array composed of three types of sensor cells expressing AgOR1, AaOR4, and AaOR15 was fabricated. Fluorescence responses were measured against solutions containing different combinations of target molecules. Specifically, solutions containing phenol, 6M5H2O, or acetophenone—either individually (single-component), in binary mixtures (two-component), or as a ternary mixture (three-component)—were applied to the array, and time-course fluorescence responses were recorded (Supplementary Fig. 8).

As shown in Fig. 6A, the multisensor array exhibited distinct fluorescence response patterns depending on the combination of target molecules in the applied solution. Comparison of the maximum fluorescence responses (Fig. 6B) revealed that the response profiles of each sensor cell type varied clearly according to the specific combination of target molecules present. Notably, the sensor cells expressing AgOR1, AaOR4, and AaOR15 maintained selective responses to phenol, 6M5H2O, and acetophenone, respectively, even in the presence of other target molecules.

**Fig. 6.**
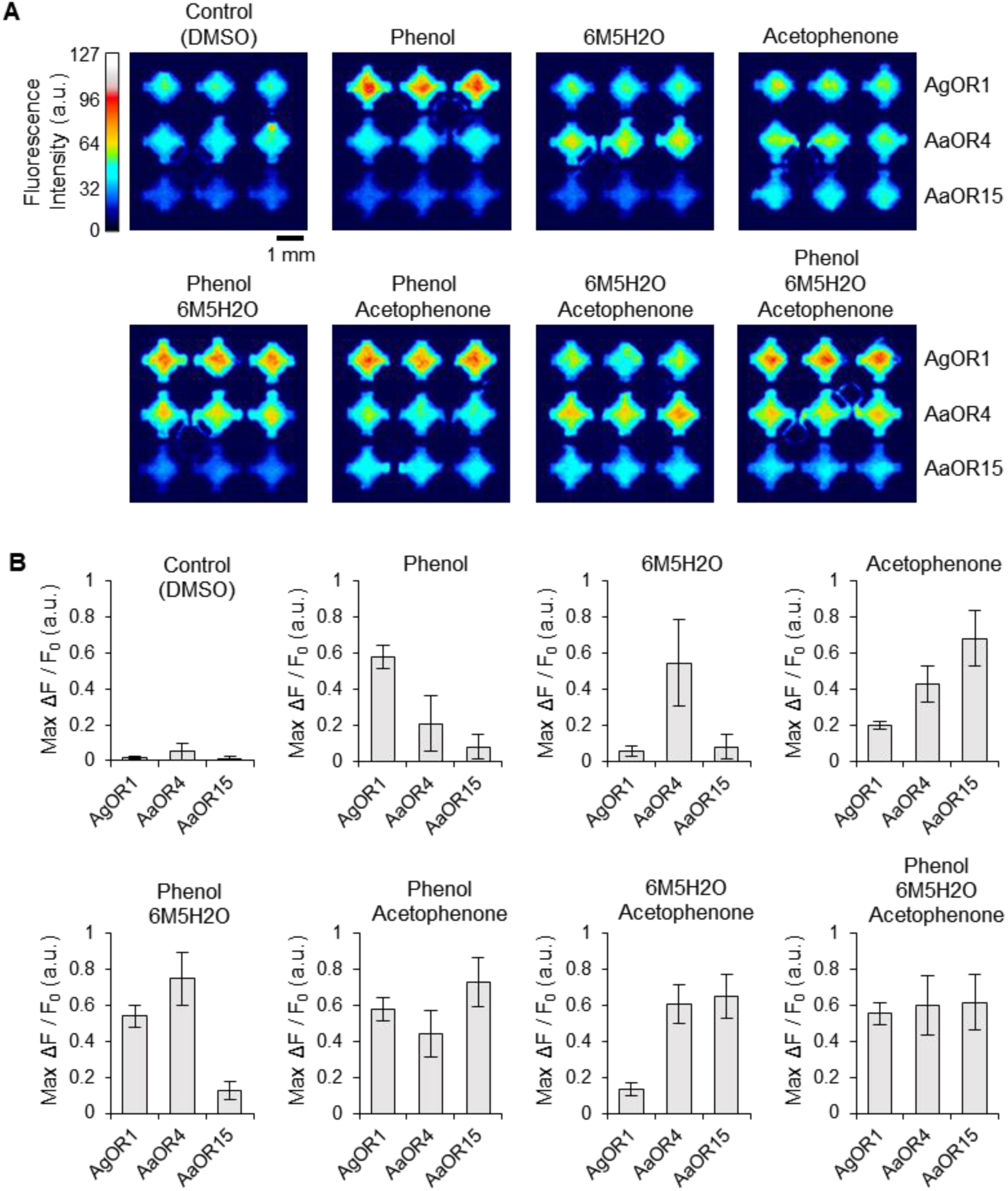
Selective and simultaneous detection of mixed target molecules using a multisensor array. (**A**) Pseudocolor fluorescence images showing the post-response of a multisensor array to aqueous solutions containing either a control (DMSO only), a single target molecule (phenol, 6-methyl-5-hepten-2-one [6M5H2O], or acetophenone), or combinations of these targets. The array comprised three types of sensor cells expressing AgOR1, AaOR4, and AaOR15. Each target molecule was used at a concentration of 100 μM, with the final DMSO concentration adjusted to 0.1% in all solutions, including mixtures. (**B**) Maximum fluorescence responses under each condition shown in (A). Each bar represents mean ± SD; *n* = 9. See also Supplementary Fig. 8 for time-course responses.

Furthermore, when multiple target molecules were present simultaneously, each of the corresponding sensor cells exhibited fluorescence responses, resulting in composite fluorescence signals that reflected the composition of the mixture. These results demonstrate that the multisensor array can not only detect single target molecules but also simultaneously identify and distinguish multiple target molecules within mixed samples. Thus, the microwell plate–based multisensor array developed in this study offers a promising approach for the simultaneous detection of target molecules in complex physiological samples.

### Detection of a target molecule added to human urine

The detection of specific biomarkers in urine offers a non-invasive and convenient approach for disease diagnosis (1-3). To detect target molecules in human urine using the developed multisensor array, we considered the potential impact of interfering substances commonly present in urine that may hinder sensor cell responses (further discussed in the Discussion section). As a solution, we devised a detection strategy involving the extraction of volatile compounds using hexane, a hydrophobic organic solvent, followed by exposure of the sensor cells to the vapor phase of the extract (Fig. 1D, E, G). In this procedure, urine samples were mixed with hexane, vortexed, and centrifuged to isolate the hexane phase containing volatile compounds from the aqueous layer (Fig. 7A). The effectiveness of this strategy was evaluated by comparing two detection approaches using urine samples containing acetophenone: (i) direct liquid-phase application and (ii) vapor-phase exposure after hexane extraction.

**Fig. 7.**
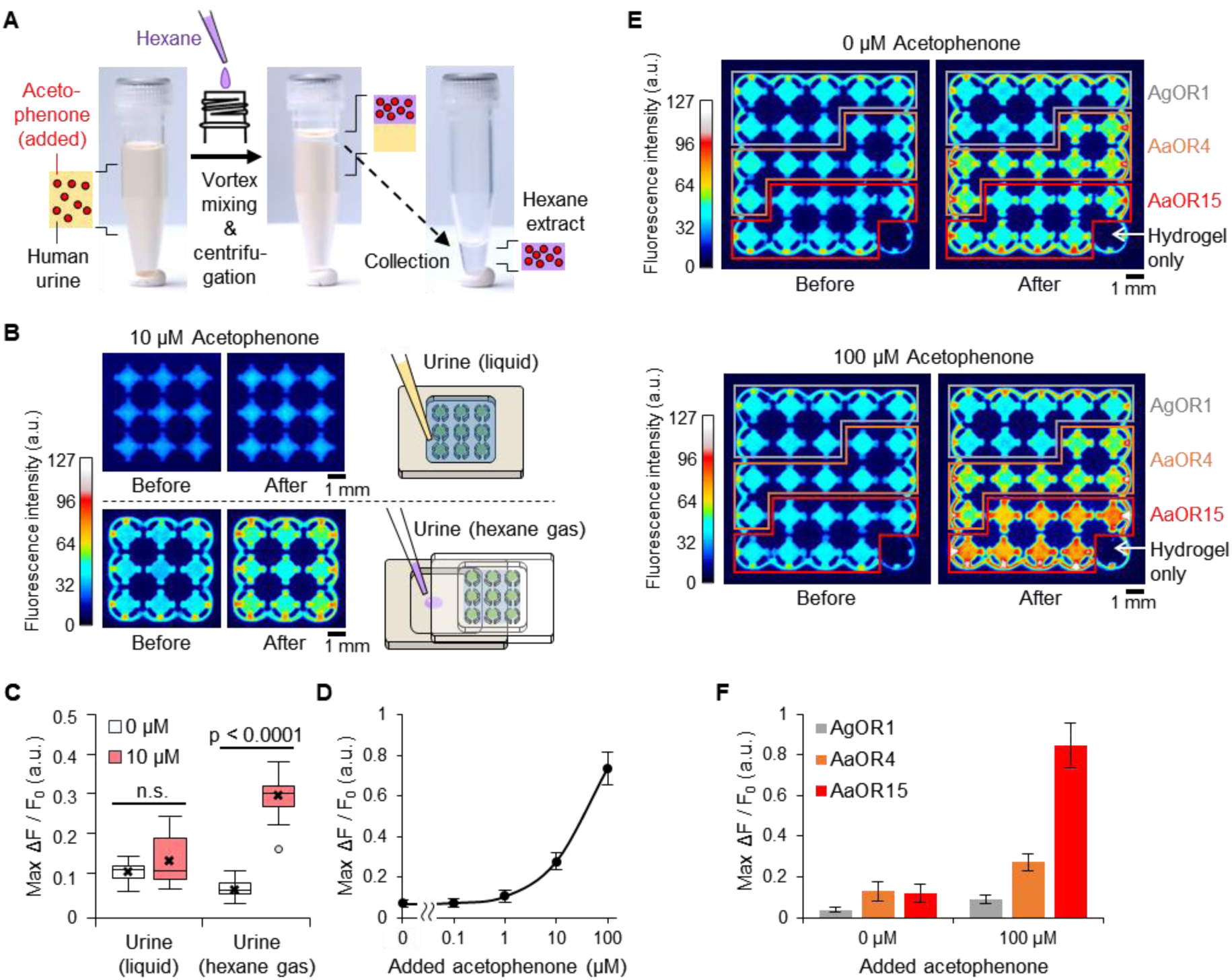
Gas-phase detection of acetophenone added to human urine using a multisensor array. (**A**) Diagram of the extraction procedure. Hexane (1:10 volume) was added to human urine, followed by vortex mixing and centrifugation. The upper hexane layer was collected as the extract. (**B**) Comparison of two detection methods using an AaOR15-expressing sensor cell array. Pseudocolor fluorescence images before and after the response are shown for direct liquid addition of urine (top) and gas-phase exposure via hexane extract (bottom). For gas-phase detection, the array was inverted after buffer removal and placed on a sample chamber. A volume of 3 μl hexane extract was added to the chamber and allowed to volatilize. (**C**) Maximum fluorescence responses of the sensor array under the two conditions shown in (B), with or without acetophenone (0 or 10 μM). Statistical analysis: Dunn’s test (liquid: *n* = 27 each; gas: 0 μM, *n* = 18; 10 μM, *n* = 27). “n.s.”, not significant. (**D**) Dose-dependent fluorescence responses of AaOR15-expressing sensor cell arrays to acetophenone extracted from urine with hexane. Note that the acetophenone concentrations indicate amounts added to the urine. Each point represents mean ± SD, *n* = 18. (**E**) Pseudocolor fluorescence images showing responses of a multisensor array to hexane extracts of urine with (100 μM; Movie 3) or without (0 μM; Movie 2) added acetophenone. The array contained AgOR1-, AaOR4-, and AaOR15-expressing cells; hydrogel-only wells showed no signal. (**F**) Maximum fluorescence responses of each sensor cell type under the conditions shown in (E). Each bar represents mean ± SD, *n* = 16. Buffers: HBSS/PIPES (pH 6.2). Excitation: intermittent illumination (2 s every ∼5 s for B–D; 2 s every ∼30 s for E–F).

As shown in Fig. 7B, AaOR15-expressing sensor cell arrays were exposed to urine samples containing 10 μM acetophenone. In method (i), the urine solution was added directly to the buffer above the array (top panel). In method (ii), most of the buffer covering the sensor cell array was removed. The array was then inverted and placed above a lower chamber containing the hexane extract (bottom panel; Fig. 1G; Fig. 2G, H), allowing exposure to volatile compounds. As a result, negligible fluorescence responses were observed with direct addition, while clear fluorescence increases were detected with vapor-phase exposure. Time-course data (Supplementary Fig. 9) also showed that no significant response occurred with 0 μM acetophenone under either condition. At 10 μM, direct liquid-phase addition still did not produce fluorescence changes distinguishable from those at 0 μM, while vapor-phase exposure induced a clear fluorescence increase. This trend was further supported by a comparison of the maximum fluorescence responses (Fig. 7C): acetophenone-containing and non-containing samples could not be distinguished under direct addition, whereas a highly significant difference (p < 0.0001, Dunn’s test) was observed under vapor-phase exposure. Additionally, the detection sensitivity for acetophenone in urine samples using hexane extraction was comparable to the concentration–response profile obtained with standard solutions (Fig. 5L), indicating that the lower limit of detection lies within the 1–10 μM range (Fig. 7D). These findings demonstrate the effectiveness of hexane extraction as a pretreatment for detecting acetophenone in human urine using the sensor cell array.

Furthermore, a multisensor array composed of AgOR1-, AaOR4-, and AaOR15-expressing sensor cells was prepared and exposed to hexane extracts of urine samples containing 0 μM (Movie 2) or 100 μM (Movie 3) acetophenone, as shown in Fig. 7E. Distinct fluorescence response patterns were observed between two conditions. As shown in Fig. 7F, comparison of the maximum fluorescence responses showed that AaOR15-expressing sensor cells exhibited the highest response to acetophenone. These results demonstrate that the developed multisensor array, when combined with appropriate sample pretreatment, enables the selective detection of target molecules in human urine samples.

## Discussion

In this study, we demonstrated the detection of a target molecule in human urine using a cell-based multisensor array constructed with a slit-integrated microwell plate. By measuring the cumulative fluorescence response from a population of sensor cells densely encapsulated in hydrogel within each microwell, we achieved improved sensitivity and reduced variability.

Furthermore, by developing a gas-phase detection system in which target molecules were extracted from urine using a volatile organic solvent and subsequently exposed to the sensor cell array, we achieved reliable detection of target molecules in urine.

### Improved sensitivity and reduced variability of fluorescence responses

Biomarker detection in physiological samples using cell-based sensors has not been previously reported. One of the major challenges in this field has been achieving sufficient detection sensitivity. Conventional approaches typically define regions of interest (ROIs) on individual cells, but the fluorescence signals from individual cells are often weak, and there is considerable variability in responses between cells, limiting both detection sensitivity and reproducibility [47]. In this study, we enlarged ROIs at the microwell level and measured cumulative fluorescence from a population of over 100,000 sensor cells immobilized within each well (Fig. 3A). In particular, we densely and three-dimensionally packed sensor cells into the confined space of each microwell, achieving an exceptionally high cell density that effectively approaches tight packing. This density far exceeds what is attainable with planar configurations, in which variability in cell seeding and detachment can easily result in uneven coverage, preventing tight cell packing. Our three-dimensional configuration led to significantly stronger fluorescence signals in each microwell. Furthermore, the high cell occupancy within each ROI suppresses signal dilution by background, contributing to substantial improvements in normalized fluorescence responses (Fig. 3B, C). As a result, the “apparent response sensitivity” at the ROI level is significantly enhanced, yielding improved detection sensitivity and reduced variability across the sensor array, even though the intrinsic sensitivity of each individual sensor cell naturally remains unchanged. This improvement in detection performance at the ROI level also has important implications when detecting trace biomarkers in real biological samples. When combined with vapor-phase detection of target molecules extracted using hexane, this approach enabled detection of a potential biomarker present in human urine (Fig. 7), as discussed below.

### Design of slit-integrated microwells for array construction

To construct the sensor cell array, we focused on two key requirements: (1) a simple and reproducible fabrication method, and (2) high response sensitivity.

Concerning the first requirement, sensor cell arraying remains a challenging process without the aid of specialized microfabrication technologies [17, 20, 37, 39, 41, 43]. To overcome this limitation in sensor cell arraying, we developed a microwell plate designed for simple and reproducible pipetting-based cell loading. The plate consists of cylindrical wells with vertically oriented sidewall slits, which play a key role in enabling stable loading of hydrogel-encapsulated sensor cells (Fig. 1B). The slits guide the pipetted material efficiently to the bottom of the well and also serve as air vents during loading, facilitating complete filling of the microwell with hydrogel. In addition, the inherent hydrophobic nature of the acrylic, the fabrication material of the microwell plate, is thought to contribute to the prevention of leakage of both aqueous and hydrogel solutions through the slits, allowing for stable retention within the well (Fig. 2C–F).

Importantly, this fabrication approach requires no specialized microfabrication technology and, with the developed microwell plate, can be performed using only standard laboratory pipetting techniques, making it accessible even to researchers without prior experience in device fabrication. Furthermore, pipetting-based fabrication is compatible with automation and scalable production using pipetting robots, thereby facilitating the practical deployment of multisensor arrays in cell-based multiplexed assays.

In addition to ease of fabrication, the slit-integrated microwell structure also fulfills the second requirement of achieving high response sensitivity. Specifically, the sidewall slits provide additional pathways for target molecules to access sensor cells within a microwell, in addition to the top opening (Fig. 1B). This design promotes multidirectional diffusion of target molecules, leading to uniform and simultaneous activation of sensor cells, which in turn improved fluorescence response sensitivity of the sensor cell arrays toward the target molecules (Fig. 3D–J).

Additionally, the partial arcs that form part of the slit-integrated microwells create clamp-like segmented structures that help secure and retain the hydrogel-encapsulated sensor cells within the wells. While primarily serving to stabilize cell placement, this structural and functional feature also played a critical role in enabling direct exposure of the sensor cells to vapor-phase target molecules, as described in the following section.

### Vapor-phase detection of extracted target molecules using sensor cell arrays

One of the major breakthroughs in this study is the finding that detecting target molecules in human urine, previously considered challenging using cell-based sensors, can be achieved through vapor-phase detection using a sensor cell array (Fig. 7). Preliminary experiments (unpublished) suggested that several factors intrinsic to urine—such as its yellow coloration, autofluorescence [64], and high concentrations of urine-specific components like urea—could interfere with the fluorescence response of sensor cells, thereby making the detection of target molecules difficult. These interfering factors are also expected to vary among individuals, raising concerns about reproducibility and general applicability in practical use.

To overcome this issue, we developed a method in which hydrophobic target molecules were extracted from urine using hexane, a hydrophobic volatile organic solvent, and 3 μL of the resulting extract was allowed to volatilize within the confined space (<1.5 ml) of the device, creating a vapor-phase environment sufficient to activate the sensor cells. This approach effectively removes water-soluble interfering substances that are not extracted into hexane, resulting in the disappearance of urine’s yellow coloration and the elimination of autofluorescence background, while enabling efficient detection of target molecules that are extracted into hexane. In practice, acetophenone in urine could not be detected when urine samples containing 10 μM acetophenone were directly applied to the sensor cell array. In contrast, when the vapor phase of hexane extracts was used for exposure, acetophenone in urine was clearly detected.

The realization of vapor-phase exposure takes advantage of two key properties of the hydrogel used to encapsulate and immobilize the sensor cells in the microwells: high moisture retention for desiccation resistance, and mechanical stability for structural support [65]. Conventionally, sensor cells must be maintained in aqueous environments to remain functional. Owing to the high water-holding capacity of the hydrogel, the cells can be exposed directly to the vapor phase without drying out. In addition, the mechanical stability of the hydrogel not only permits the removal of buffer to expose the cells directly to vapor but also preserves the structural integrity of the sensor cell array even when the microwell device is inverted. This orientation enables a “hanging configuration,” in which volatilized target molecules are uniformly and efficiently delivered to the sensor cells.

The experiment was conducted using only 1 mL of urine and 0.1 mL of hexane for extraction. Only a few microliters of the hexane extract were introduced into the device, and the extract was fully volatilized during measurement. Future improvements in sensitivity may be achieved by increasing the volume of urine used for extraction to enrich the concentration of target molecules, or by optimizing the choice of volatile organic solvents based on the physicochemical properties of specific targets. These findings support the development of a practical and scalable strategy for noninvasive detection of cancer-related volatile biomarkers in urine using sensor cell arrays.

### Current limitations and future perspective

Expanding the library of sensor cells expressing olfactory receptors (ORs) will be critical for detecting a broader range of biomarker candidates. Insect-derived ORs, known for excellent molecular recognition and broad responsiveness [54, 56], are attractive for this purpose.

Combining multiple OR-expressing sensor cells could enable detection and discrimination of complex target mixtures.

Although we successfully detected 10 μM acetophenone in human urine, enhancing sensitivity to detect lower concentrations remains a key challenge. While our approach effectively harnessed the responsiveness of sensor cells, optimization of the sensor cells themselves is needed. Stable OR-expressing lines were used here, but cells were not selected based on their fluorescence response to target molecules. Variability among individual cells may still limit overall detection sensitivity. Future efforts to isolate highly expressing and responsive cells using cell sorters may yield more homogeneous populations, enabling the development of even more sensitive cell-based sensors.

The method developed here provides a promising platform for diagnostic applications using cell-based sensors. Its utility could extend beyond urine to other biological fluids, such as blood, saliva, and tear fluid. The sensor cell array, comprising a compact microwell plate and hydrogel-encapsulated cells, enables gas-phase detection with minimal sample volumes. This structural and functional design supports the development of simple, portable, on-site diagnostic systems. In the future, this platform may serve as a foundation for advancing non-invasive biomarker screening technologies for early detection of cancers, infectious diseases, and metabolic disorders.

Taken together, this study establishes a strategy for constructing a cell-based multisensor array using a slit-integrated microwell plate and densely packed sensor cells, enabling gas-phase detection of volatile target molecules extracted from urine. These findings highlight the potential for expanding the application range of cell-based sensors in biomedical diagnostics.

## Methods

### Materials and reagents

Poly(methyl methacrylate) (PMMA) substrates were purchased from Mitsubishi Chemical (Japan). TrueGel3D (TRUE7) was obtained from Sigma-Aldrich (USA). Acetophenone, phenol, 6-methyl-5-hepten-2-one (6M5H2O), and hexane were purchased from Kanto Chemical (Japan). ExpiSf9 cells and ExpiSf CD Medium were obtained from Thermo Fisher Scientific (USA). G-418 and penicillin-streptomycin solution (×100) were purchased from FUJIFILM Wako Pure Chemical Corporation (Japan). Puromycin was purchased from Invivogen (USA). Hanks’ Balanced Salt Solution (HBSS) (+) with Ca and Mg, without phenol red, was obtained from Nacalai Tesque (Japan). HEPES and PIPES were purchased from Dojindo Laboratories (Japan). All other chemicals were obtained from FUJIFILM Wako Pure Chemical Corporation (Japan).

All reagents were used without further purification. All aqueous solutions were prepared using ultrapure water from a Milli-Q system (Milli-Q Integral 3, Merck, Germany). For coloring Milli-Q water, watercolor paint was used. Sodium hypochlorite bleach (product name: Haiter) was purchased from Kao Corporation (Japan). Pooled urine from healthy donors was obtained from Medix Biochemica (USA).

### Design and fabrication of microwell plates

Microwell plates (24 mm × 36 mm, 4 or 5 mm in height) in 3 × 3 (9 microwells), 5 × 5 (25 microwells), 7 × 7 (49 microwells), and 9 × 9 (81 microwells) formats were fabricated using transparent or black PMMA substrates (Supplementary Fig. 2). Each microwell had a cylindrical shape formed by partial arcs, with an inner diameter of 1.2 mm, an outer diameter of 1.6 mm, and a height of 1.0 mm. The sidewalls, 0.2 mm thick, were integrated with vertical slits defined by a central angle (θ). The microwells were placed inside a chamber with a depth of 3 mm or 4 mm and arranged at 0.4 mm intervals, starting 2.0 mm from the inner wall of the chamber. The microwell plates were designed using CAM software (Alphacam, Vero Software, UK), and micromachined from 4- or 5-mm-thick PMMA substrates using a milling machine (MM-100, Modia Systems, Japan). Before reuse in experiments, the microwell plates were immersed in approximately 10-fold diluted sodium hypochlorite bleach solution (final concentration of sodium hypochlorite <1%), followed by ultrasonic cleaning.

### Generation of sensor cells

ExpiSf9 cells were seeded at a density of 6 × 10⁵ cells per well in a 6-well plate using ExpiSf CD medium. A PiggyBac transposon vector encoding the jGCaMP7s gene and a neomycin resistance gene, along with mRNA encoding the transposase gene, were transfected into the cells at 1 μg each using the TransIT-Insect Transfection Reagent (Takara Bio, Japan), according to the manufacturer’s protocol. At 3 and 6 days post-transfection, the culture medium was replaced with ExpiSf CD medium supplemented with G418 at final concentrations of 125 μg/mL and 250 μg/mL, respectively. On day 9 post-transfection, cells were detached by pipetting and transferred to a 125 mL Erlenmeyer flask (Thermo Fisher Scientific, USA) containing 16 mL of ExpiSf CD medium supplemented with 250 μg/mL G418. The cells were cultured at 27°C with shaking at 125 rpm to promote proliferation, resulting in the establishment of a stable cell line expressing jGCaMP7s. Subsequently, the established jGCaMP7s-expressing cell line was further transfected with a PiggyBac transposon vector encoding three genes: the OR gene of interest, a high-sensitivity chimeric Orco gene [66], and a puromycin resistance gene. Transfections were carried out alongside mRNA encoding the transposase gene, following the same protocol. Stable cell lines were selected using puromycin at sequential concentrations of 2.5 μg/mL and 5 μg/mL. The final selected cells (sensor cells) were used for all subsequent experiments.

### Culture of sensor cells

Sensor cells derived from the insect cell line ExpiSf9 were cultured in non-baffled shake flasks at 27°C with shaking at 125 rpm. ExpiSf CD medium supplemented with penicillin-streptomycin solution (×1) was used for cell culture. Cells were passaged every 3–4 days by diluting them to a density of 0.5×10⁶ cells/mL. For sensor cell array fabrication, cells cultured for 2–4 days after passage were used.

### Encapsulation of sensor cells in hydrogel and arraying in microwell plates

Sensor cells suspended in culture medium were pelleted by centrifugation at 300 × g for 5 minutes using a swing rotor (T5SS31, Himac, Japan), and the supernatant containing the culture medium was carefully removed. The collected cells were resuspended in TrueGel3D (TRUE7), prepared according to the manufacturer’s protocol except for the crosslinker, at a final concentration of 1×10⁸ cells/mL. The crosslinker was then added to the cell suspension, followed by gentle mixing. Subsequently, 1.1 μL of the suspension was dispensed into each microwell using the multi-dispensing mode of an electronic pipette (Xplorer, Eppendorf, Germany). After dispensing, the microwell plates were incubated at 27 °C for 25 minutes to allow complete gelation of the hydrogel containing the sensor cells. After gelation, the chamber of each microwell plate was filled with HBSS containing 0.1% (w/v) bovine serum albumin (BSA), buffered with either 20 mM HEPES (pH 7.2) or 20 mM PIPES (pH 6.2), referred to as HBSS/HEPES/BSA or HBSS/PIPES/BSA, respectively. The volume of buffer (HBSS/HEPES/BSA or HBSS/PIPES/BSA) added to the chamber was adjusted according to the format of the microwell plate: 200 μL for 3 × 3, 400 μL for 5 × 5, 600 μL for 7 × 7, and 1000 μL for 9 × 9 formats. Microwell plates with sensor cell arrays were stored at 4 °C until use and equilibrated to room temperature prior to target molecule detection.

### Microscopic observation of sensor cells immobilized in microwells

Sensor cells immobilized in microwells through hydrogel encapsulation were observed using a 55× zoom stereomicroscope (MS-2502T, BioTools, Japan) equipped with a fluorescence illumination unit (BT-ExSMH-BG, BioTools, Japan), which was configured with a blue excitation light (485 nm) and a green emission filter (530 nm). Both bright-field and fluorescence images were acquired.

### Fluorescence response measurement of sensor cell arrays

Fluorescence imaging of sensor cell arrays prepared using microwell plates was performed using a fluorescence imaging system (DP-T130z, BioTools, Japan) equipped with an 8-bit (256 grayscale) CMOS camera, a blue excitation light (485 nm), and a green emission filter (530 nm). The sensor cell arrays were placed inside the imaging box of the system. Imaging was carried out at room temperature under continuous blue light illumination or intermittent illumination for approximately 1–2 seconds in each 5-second cycle. The blue light was turned on and off either manually or using an interval timer (FT-022, Tokyo Garasu Kikai, Japan).

After the start of image acquisition, the door of the imaging box was opened, and a 5× concentrated aqueous solution of the target molecule (final concentrations: 0–100 μM) was added to the chamber of the microwell plate by pipetting. The stock solution of each target molecule was first prepared in dimethyl sulfoxide (DMSO), then diluted with buffer (HBSS/HEPES or HBSS/PIPES) to prepare the 5× working solution. The buffer used for dilution had the same pH as that in the chamber but did not contain BSA. The final DMSO concentration was adjusted to not exceed 0.3% (v/v). The volume of the target solution added to the chamber was adjusted according to the microwell plate format: 50 μL for 3 × 3, 100 μL for 5 × 5, 150 μL for 7 × 7, and 250 μL for 9 × 9 formats. The addition was performed either in a single step or in four sequential dispenses using the multi-dispensing mode of an electronic pipette (Xplorer, Eppendorf, Germany).

For image analysis, ImageJ software (version 1.52a, NIH, USA) was used. Circular regions of interest (ROIs) were defined within each microwell, and the average fluorescence intensity was quantified. The fluorescence response of sensor cells was calculated as the normalized change in fluorescence intensity (ΔF/F₀), where ΔF = F − F₀. Here, F is the fluorescence intensity at each time point, and F₀ is the baseline intensity immediately before the response—i.e., just after closing the imaging box following the addition of the target molecule (approximately 40 seconds after the start of recording). The maximum value of the normalized fluorescence change (Max ΔF/F₀) was extracted as the peak response following the addition of the target molecule. The fluorescence response was plotted using the time of target addition (typically 30 seconds after the start of recording) as time zero (0 s) to facilitate direct comparison of the cellular responses across assays.

### Hexane extraction of acetophenone added to human urine

Acetophenone was first dissolved in dimethyl sulfoxide (DMSO) and then added to pooled urine samples derived from healthy donors to achieve final concentrations ranging from 0 to 100 *μ*M. The final concentration of DMSO was adjusted to 0.1% (v/v). The urine samples containing acetophenone were transferred to screw-cap microtubes, and hexane was added at a urine-to-hexane volume ratio of 10:1 (Fig. 7A). The mixture was vigorously vortexed for 20 minutes and then centrifuged at 21,600 × g for 5 minutes using an angled rotor (T15A44, Himac, Japan), resulting in the separation of an upper (hexane) phase and a lower (urine-derived aqueous) phase. The hexane phase was carefully transferred to a separate screw-cap microtube and stored at – 80 °C until use.

### Gas-phase detection of hexane extracts using sensor cell arrays

Sensor cell arrays were prepared using transparent microwell plates filled with HBSS/PIPES/BSA solution, as described in “Encapsulation of sensor cells in hydrogel and arraying in microwell plates.” Just before use, 90% of the solution in the chamber was removed (Fig. 1D). A small container made of PMMA, with the same width and depth as the microwell plate, was placed inside the imaging box of the fluorescence imaging system. This container included a sample chamber measuring 21.6 mm × 21.6 mm in area and 3 mm in height. The microwell plate was then inverted and placed on top of the container such that the side with the sensor cell array faced the sample chamber (Fig. 2G, H). At this stage, the microwell plate and the container were slightly offset to create a gap, allowing the hexane extract to be dispensed into the chamber.

Fluorescence imaging and analysis were performed similarly to the procedure described in “Fluorescence response measurement of sensor cell arrays,” with specific modifications as outlined below. Imaging was conducted at room temperature under intermittent blue light illumination: either 2 seconds in each 5-second cycle or 2 seconds in each 30-second cycle. After the start of image acquisition, the door of the imaging box was opened, and 3 μL of the hexane extract was pipetted into the sample chamber. The offset between the microwell plate and the container was promptly corrected, thereby sealing the sample chamber with the sensor cell array positioned above it. This setup allowed the target molecules in the hexane extract to volatilize within the enclosed small space (<1.5 mL), thereby exposing the sensor cells to gas-phase molecules. During the imaging period, the entire volume of the hexane extract volatilized.

For comparison, a control experiment was also conducted by directly adding 50 μL of a urine sample containing 10 μM acetophenone into 200 μL of buffer covering the sensor cell array. Fluorescence imaging was performed in the same manner as described above.

### Statistical analysis

Statistical analysis was performed only for Fig. 7C, using the Kruskal–Wallis test followed by post hoc comparisons with Dunn’s test and Bonferroni correction. All analyses were conducted using the R statistical software (http://www.R-project.org). All bar graphs represent mean values ± standard deviation (SD).

### Data availability

Data supporting the results of this study are available from the corresponding authors upon request.

## Acknowledgments

We thank T. Iida and M. Kiji (KISTEC) for their technical support during the experiments.

## Funding

This work was supported by JST CREST JPMJCR20C4 (S.Takeuchi, T.O., and Y.T.) and JSPS KAKENHI 23K06733 (H.M.).

### Author contributions

Conceptualization: S.Takeuchi, T.O., Y.T., H.O., H.M. Methodology: H.M., Y.K., N.S., T.O., H.O., S.Takamori, S.Takeuchi Investigation: H.M., Y.K., N.S.

Visualization: H.M.

Supervision: S.Takeuchi, T.O., Y.T. Writing—original draft: H.M., N.S.

Writing—review & editing: S.Takeuchi, T.O., H.O., S.Takamori, N.S., Y.K., Y.T., H.M.

## Data and materials availability

All data needed to evaluate the conclusions in the paper are available in the main text or the Supplementary Materials.

## Supplementary Materials

Please refer to the separately submitted Supplementary Materials for additional information, including Supplementary Figures S1–S9 and Movies 1 –3.

## Supplementary Information

This file includes:

Supplementary Figs. 1 to 9

Legends for Supplementary Movies 1 to 3

---

**Supplementary Fig. 1.**
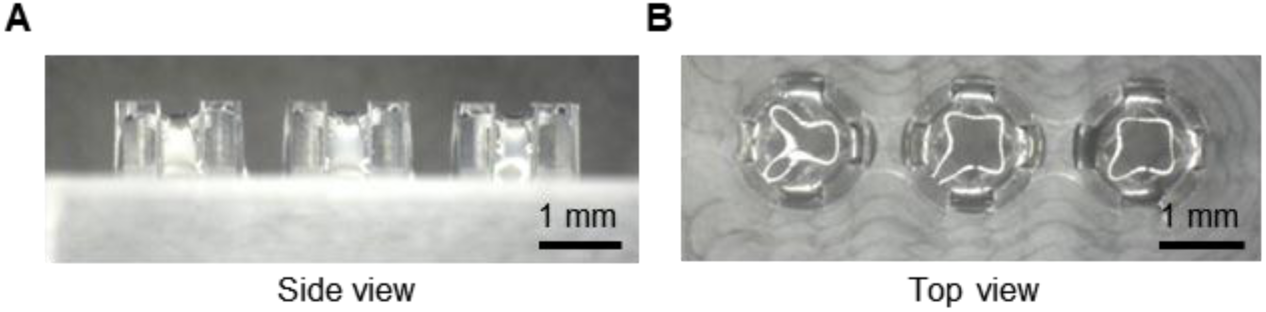
Retention of aqueous solution in slit-integrated microwells. (**A**, **B**) Side (A) and top (B) views of slit-integrated microwells retaining a water droplet.

**Supplementary Fig. 2.**
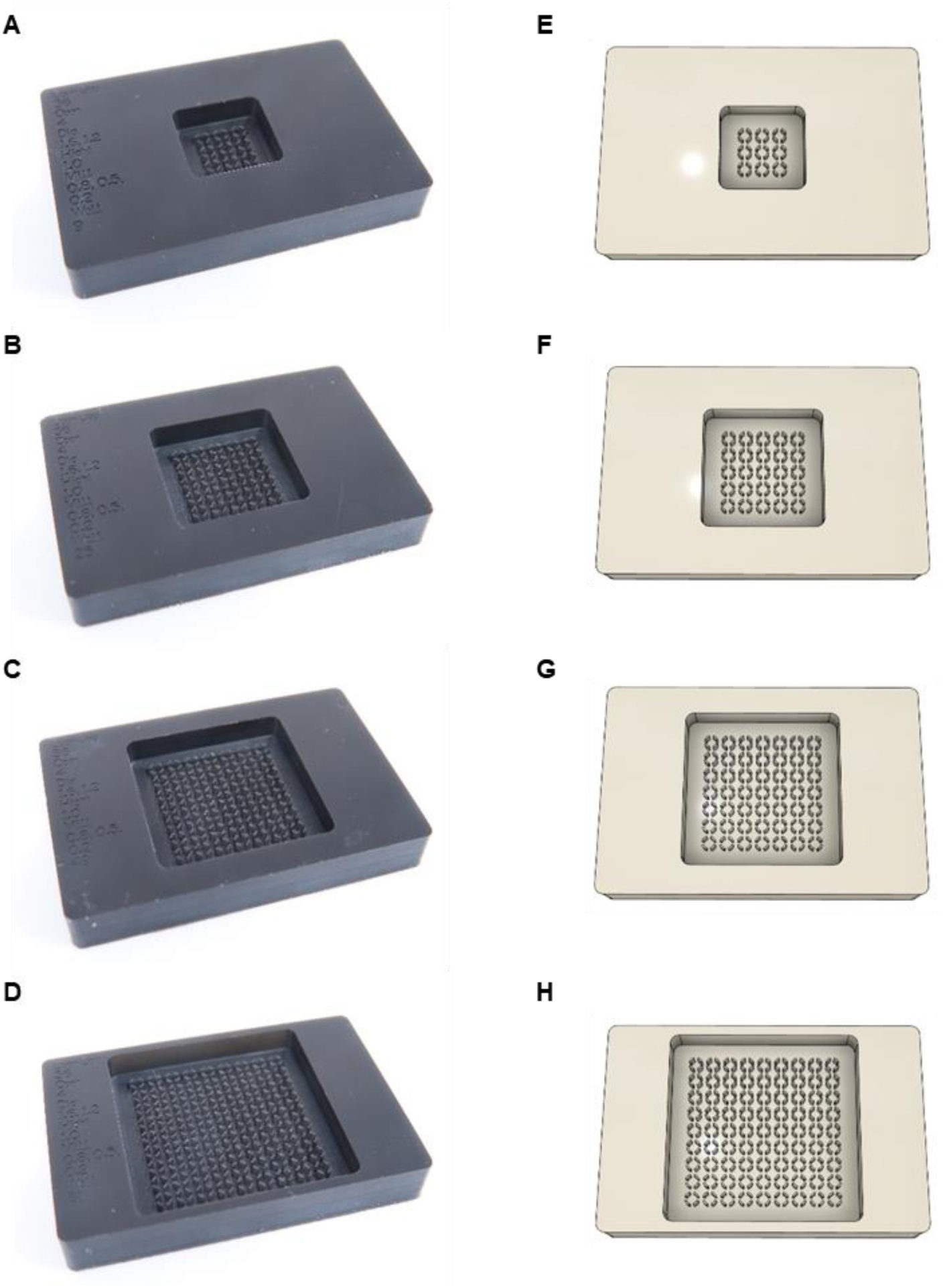
Design of slit-integrated microwell plates. (**A**, **E**) 3×3 format (9 microwells); (**B**, **F**) 5×5 format (25 microwells); (**C**, **G**) 7×7 format (49 microwells); (**D**, **H**) 9×9 format (81 microwells). (**A**–**D**) Photographs of slit-integrated microwell plates fabricated from black acrylic. (**E**–**H**) Schematic illustrations corresponding to the microwell formats shown in (**A**–**D**).

**Supplementary Fig. 3.**
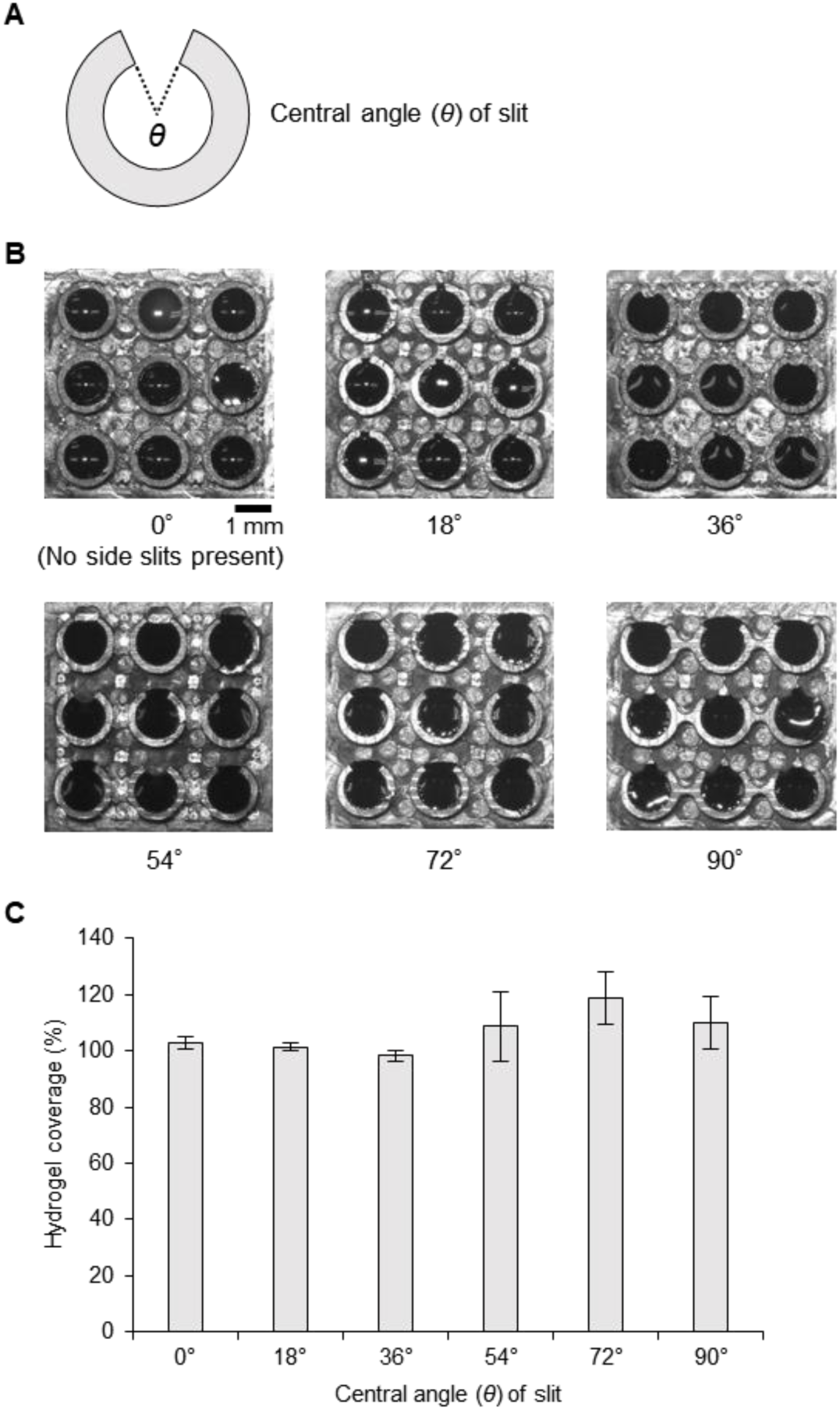
Retention of hydrogel-encapsulated sensor cells in microwells with no slit or a single slit. (**A**) Schematic illustration of a microwell with a slit defined by a central angle (θ). (**B**) Stereomicroscope images (M80, Leica, Germany) of hydrogel-encapsulated sensor cells retained in 3×3-format microwell plates composed of microwells with no slit (θ = 0°) or a single slit with varying central angles (18°, 36°, 54°, 72°, or 90°). The hydrogel appears dark due to the black acrylic background of the microwell plate. (**C**) Hydrogel coverage (%) in each microwell, calculated as the area ratio of the hydrogel (dark region) to the inner area of the microwell (1.18 ± 0.01 mm²; mean ± SD). The inner area was measured using a digital microscope (VHX-1000, KEYENCE, Japan), and the hydrogel area was quantified from the images in (B) using the Freehand Selections tool in ImageJ (v1.52a, NIH, USA). Each bar represents the mean ± SD of *n* = 9 microwells.

**Supplementary Fig. 4.**
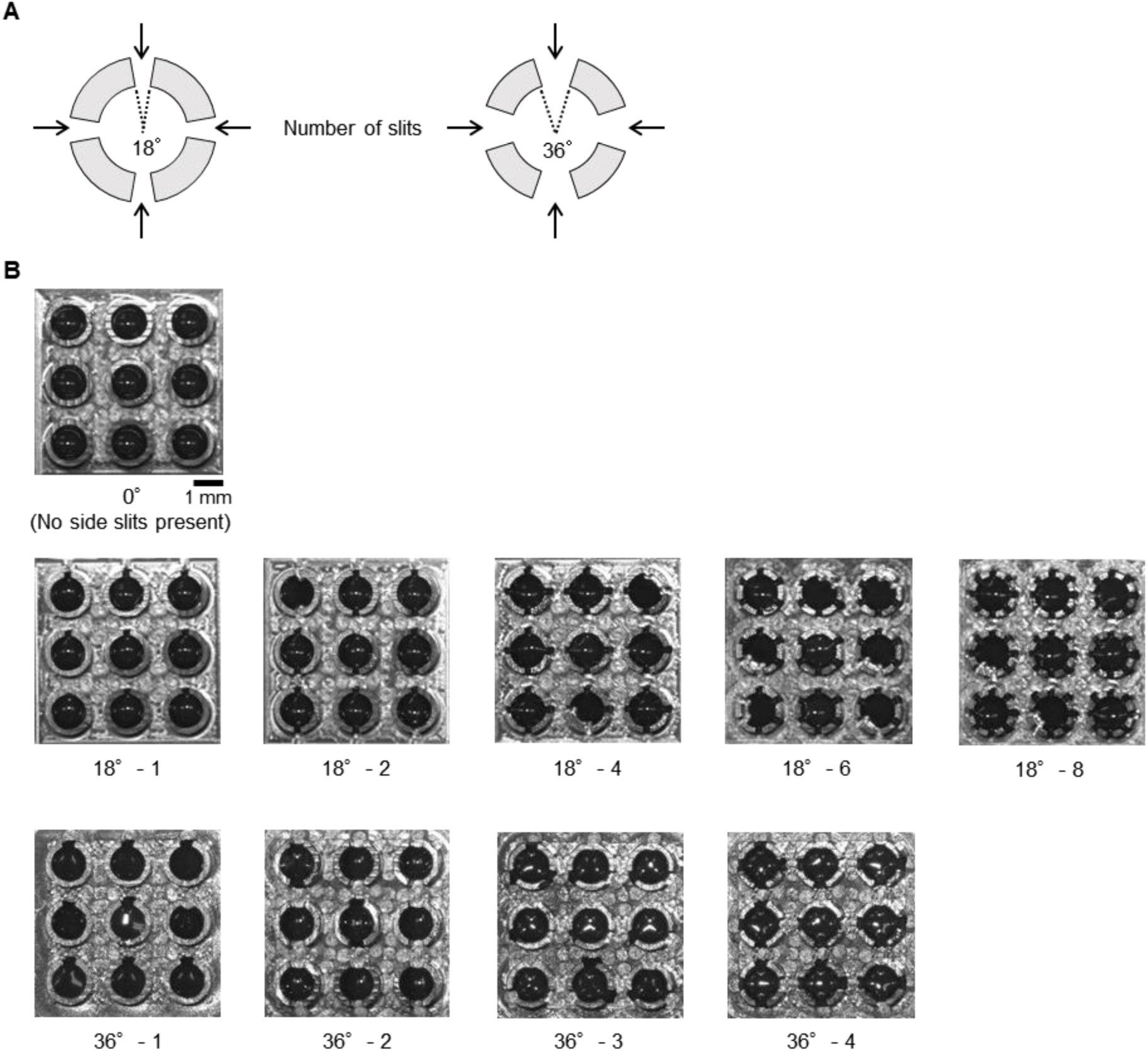
Slit-integrated microwells with multiple slits for retaining hydrogel-encapsulated sensor cells. (**A**) Schematic illustrations of microwells with four slits, each slit having a central angle of either 18° or 36°. (**B**) Stereomicroscope images of 3×3-format microwell plates composed of microwells with no slits (0°), or with multiple slits having central angles of 18° (1–8 slits) or 36° (1–4 slits). The microwells were filled with hydrogel-encapsulated sensor cells. Images were acquired using the same method as described in fig. S3.

**Supplementary Fig. 5.**
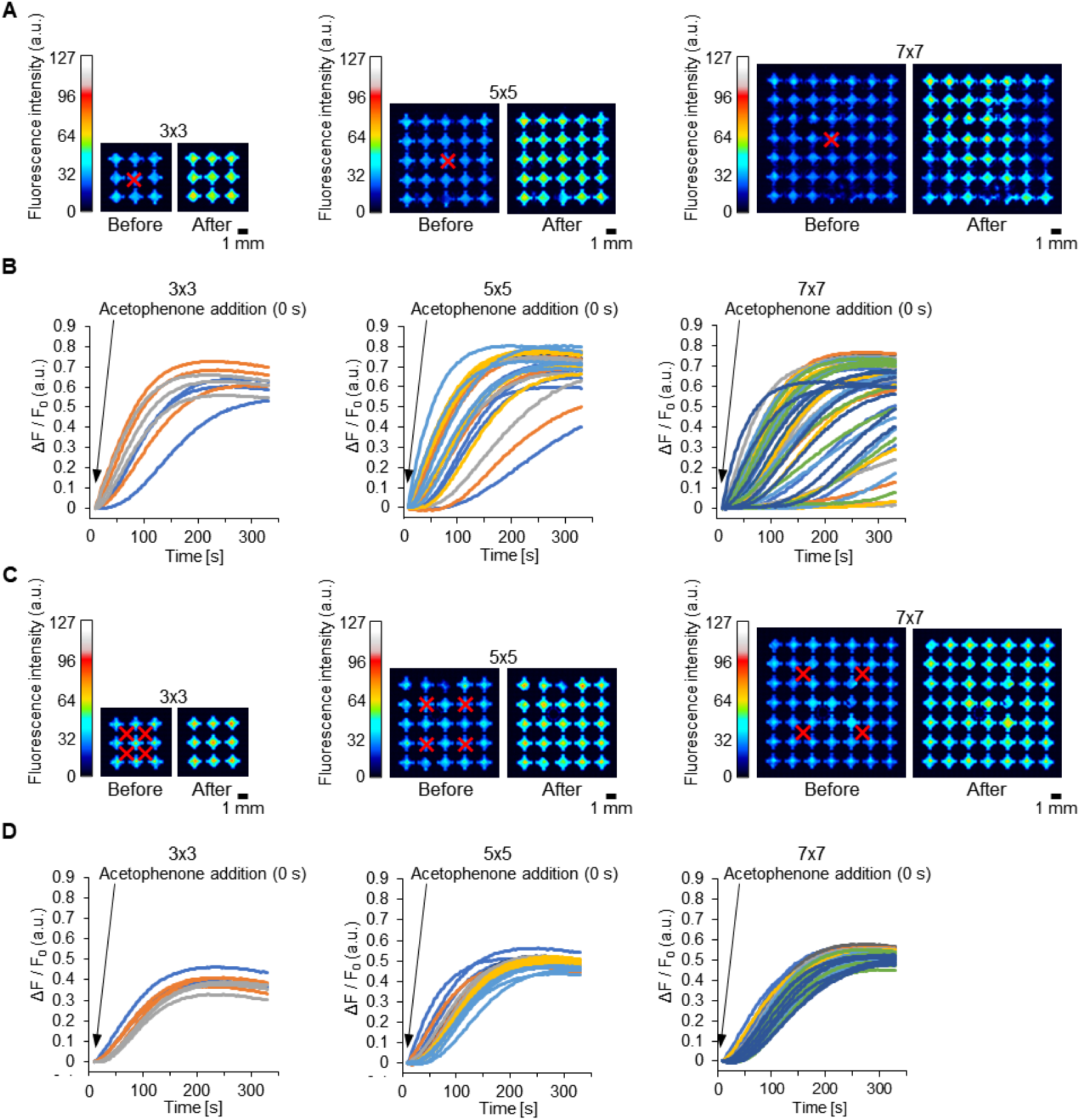
Effect of the number of target molecule additions on the consistency of sensor cell responses across different microwell plate formats. (**A**, **C**) Pseudocolor fluorescence images showing responses of sensor cell arrays (expressing AaOR15) in slit-integrated 3×3, 5×5, and 7×7 microwell plates, with each microwell containing four slits (central angle: 36°). Red crosses indicate the locations where acetophenone solution (100 µM final) was added either in a single step (A) or in four separate steps (C). Before: pre-response (12 s after addition); After: post-response (300 s after addition). (**B**, **D**) Time-course fluorescence responses corresponding to the conditions in (A) and (C), respectively: single addition (B), four additions (D). Line colors indicate microwell plate rows: blue (row 1), orange (2), gray (3), yellow (4), light blue (5), green (6), dark blue (7). Assay buffer and total sample volumes were as follows: 3×3 format—200 µL buffer, 50 µL sample; 5×5 format—400 µL, 100 µL; 7×7 format—600 µL, 150 µL.

**Supplementary Fig. 6.**
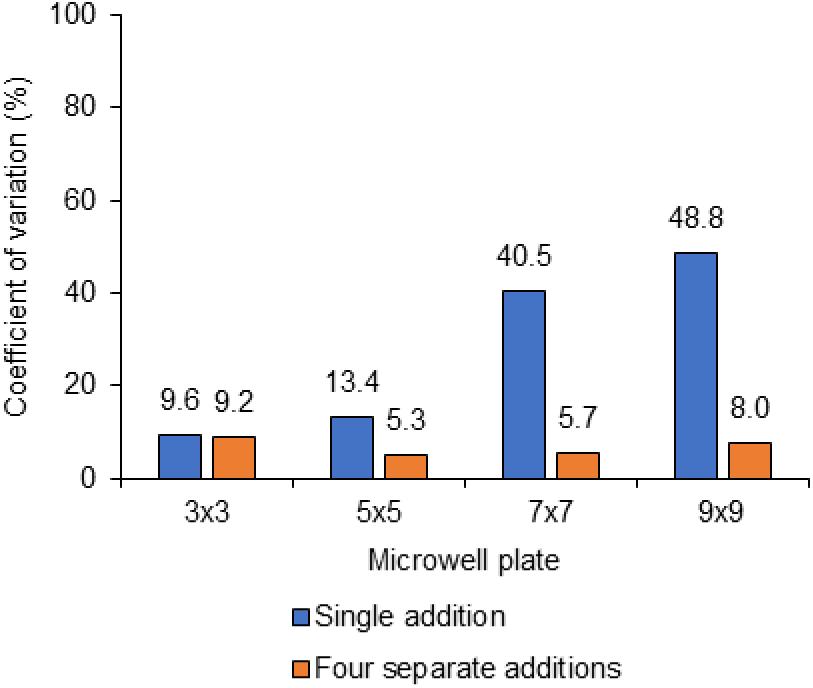
Comparison of fluorescence response variability under different target molecule addition protocols. The coefficient of variation (CV) was calculated from the maximum normalized fluorescence response (Max ΔF/F₀) of sensor cells subjected to either a single addition or four separate additions of the target molecule. Sensor cells were arrayed in microwell formats of 3×3, 5×5, 7×7, and 9×9, with 9, 25, 49, and 81 microwells analyzed for each format, respectively. CV values were derived from Fig. 4C, D (9×9 format) and fig. S5B, D (3×3, 5×5, and 7×7 formats). CV values are shown above each bar; blue bars represent single additions, and orange bars represent four separate additions.

**Supplementary Fig. 7.**
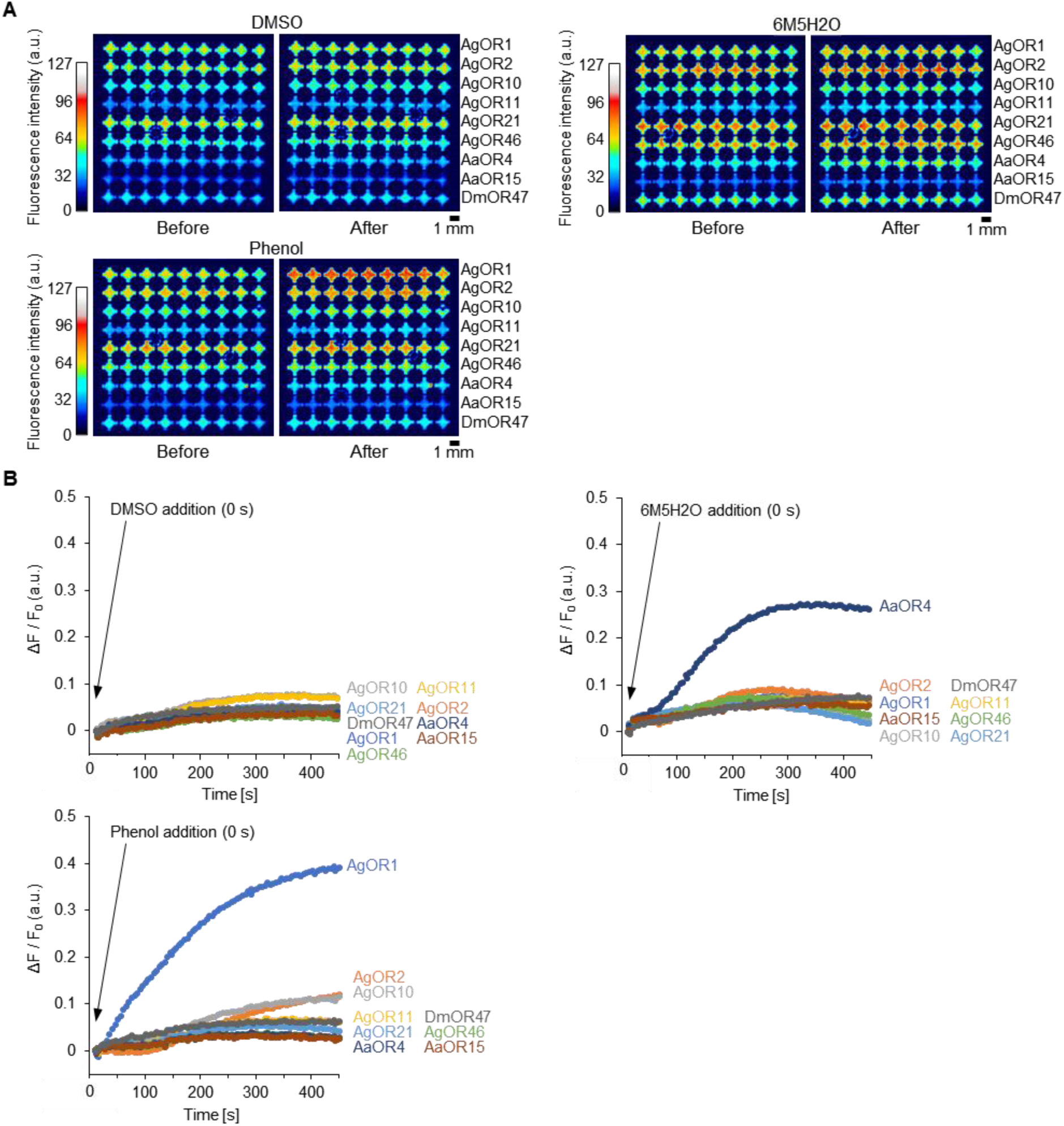
Fluorescence responses of a multisensor array to target molecules and a negative control. (**A**) Pseudocolor fluorescence images showing responses of a multisensor array composed of nine sensor cell types, each expressing a different olfactory receptor (OR) and arranged by type in horizontal rows. Either control (DMSO only), phenol, or 6-methyl-5-hepten-2-one (6M5H2O) was added in four separate steps (final concentration: 100 µM for phenol and 6M5H2O). All solutions contained 0.1% DMSO. Before: pre-response (10–12 s after addition); After: post-response (401 s after addition). (**B**) Time-course fluorescence responses under the same conditions as in (A). Each curve represents the mean response from *n* = 9 microwells.

**Supplementary Fig. 8.**
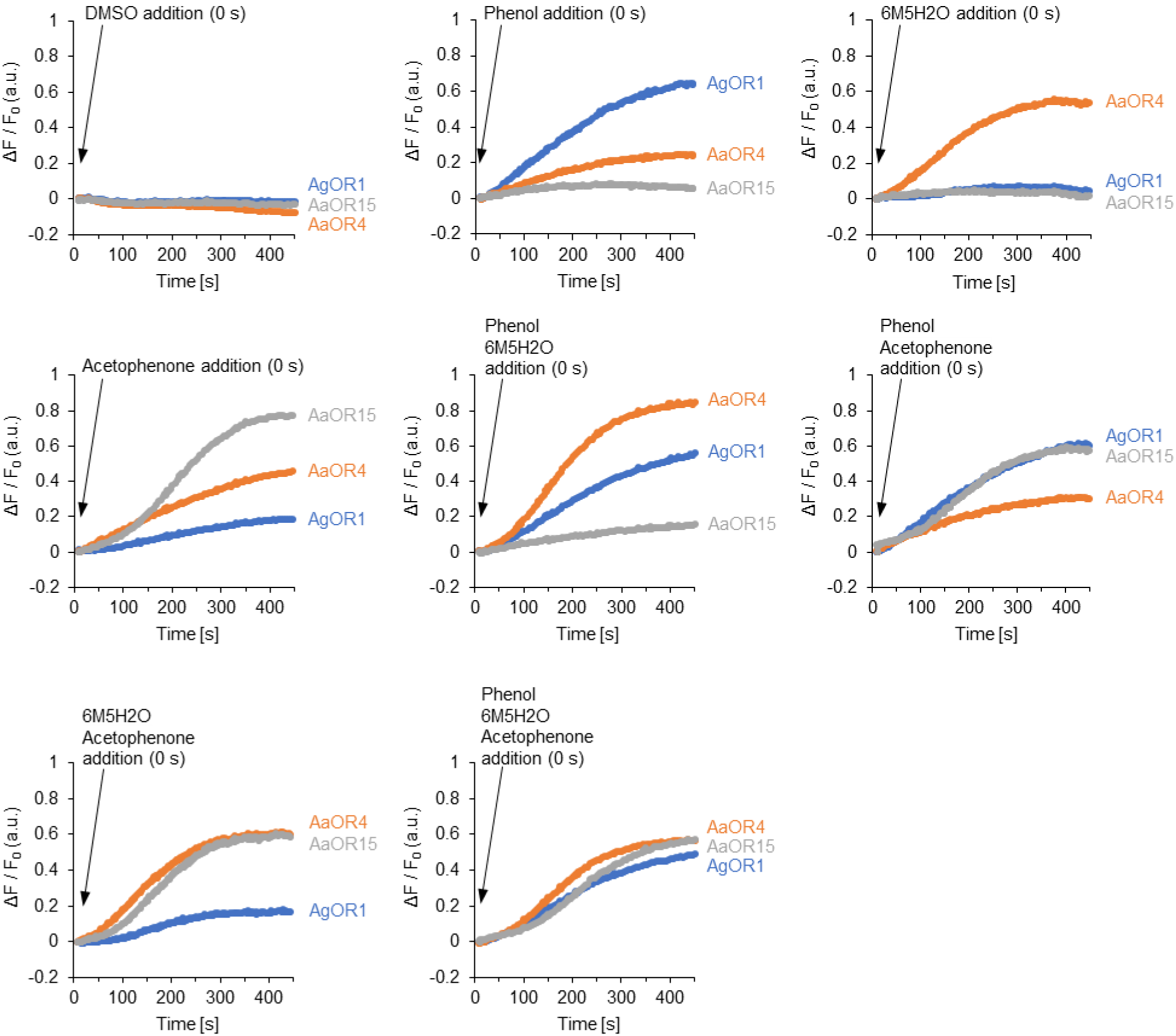
Time-course fluorescence responses for the selective and simultaneous detection of multiple target molecules using a multisensor array. Time-course fluorescence responses measured under the same conditions as in Fig. 6. Aqueous solutions containing either control (DMSO, no target molecule), phenol, 6-methyl-5-hepten-2-one (6M5H2O), acetophenone, or their mixtures were added to the array. The array consisted of three sensor cell types expressing AgOR1, AaOR4, and AaOR15. For each cell type, the average response from three microwells is shown. Final concentrations: 100 µM for each target molecule; all solutions contained 0.1% DMSO.

**Supplementary Fig. 9.**
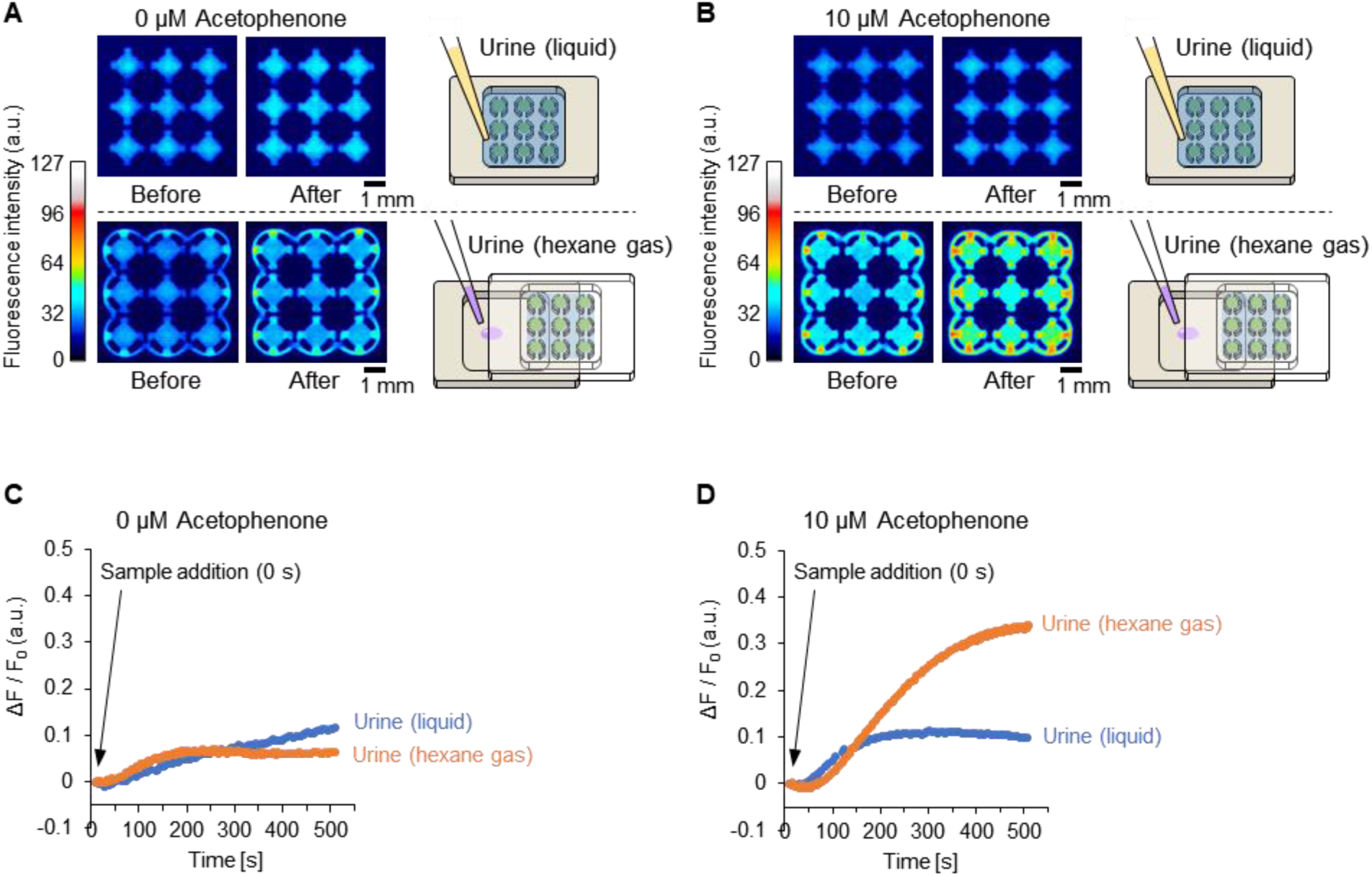
Comparison of liquid-phase and gas-phase detection of acetophenone added to human urine using a sensor cell array. (**A**) Pseudocolor fluorescence images comparing the responses of a sensor cell array to human urine without added acetophenone. Top: urine was directly added to the array filled with buffer. Bottom: a hexane extract of the urine sample was added to a sample chamber, exposing the cells to volatilized compounds in the confined space (< 1.5 mL) between the chamber and the inverted array (after buffer removal). (**B**) Pseudocolor fluorescence images comparing responses to urine samples containing 10 µM acetophenone. Experimental procedures were the same as in (A). Top: direct addition; bottom: gas-phase exposure via hexane extract. This panel is identical to Fig. 7B. (**C**, **D**) Time-course fluorescence responses corresponding to the conditions in (A) and (B), respectively. Each curve represents the mean of *n* = 9 microwells. Acetophenone concentrations indicate the amounts added to the urine samples.

**Movie 1. Continuous pipetting of water into slit-integrated microwells using an electric pipette.**

Time-lapse video showing sequential dispensing of Milli-Q water into a slit-integrated microwell plate using an electric pipette. Stable droplets were rapidly formed and retained in the microwells without spreading or merging. This demonstrates the feasibility of rapid and reproducible array formation via simple pipetting.

**Movie 2. Fluorescence response of a multisensor array to human urine without acetophenone (gas-phase exposure).**

Time-lapse pseudocolor fluorescence imaging of a multisensor array composed of AgOR1-, AaOR4-, and AaOR15-expressing sensor cells. The array was inverted and placed over a chamber, into which a hexane extract of human urine without added acetophenone (0 µM) was introduced. No significant fluorescence response was observed. This video corresponds to Fig. 7E (0 µM condition).

**Movie 3. Fluorescence response of a multisensor array to acetophenone (100 µM) in human urine (gas-phase exposure).**

Time-lapse pseudocolor fluorescence imaging of a multisensor array composed of AgOR1-, AaOR4-, and AaOR15-expressing sensor cells. The array was inverted and placed over a chamber, into which a hexane extract of human urine containing acetophenone (100 µM) was introduced. As acetophenone volatilized, distinct fluorescence responses were observed, particularly in AaOR15-expressing cells. This video corresponds to Fig. 7E (100 µM condition).

